# Stromal STAT5-mediated trophic activity regulates hematopoietic multipotent progenitor niche factors

**DOI:** 10.1101/2022.10.07.511255

**Authors:** Zhengqi Wang, Grace Emmel, Hong Seo Lim, Wandi Zhu, Astrid Kosters, Eliver E.B. Ghosn, Peng Qiu, Kevin D. Bunting

## Abstract

Signal transducer and activator of transcription 5 (STAT5a and STAT5b) are intrinsically critical for normal hematopoiesis but are also expressed in stromal cells. Here, STAT5ab knockout (KO) was generated with a variety of bone marrow hematopoietic and stromal Cre transgenic mouse strains. Vav1-Cre, the positive control for loss of multipotent hematopoietic function, surprisingly dysregulated niche factor mRNA expression and deleted STAT5ab in CD45^neg^ cells. Single cell transcriptome analysis of bone marrow from wild-type or Vav1-Cre KO mice showed hematopoietic stem cell myeloid commitment priming and upregulated protein translation genes. Nes^+^ cells were detected in both CD45^neg^ and CD45^+^ clusters and deletion of STAT5ab with Nes-Cre caused hematopoietic repopulating defects. To follow up on these promiscuous Cre promoter deletions in CD45^neg^ and CD45^+^ bone marrow cell populations, more stroma-specific Cre strains were generated and demonstrated reduction in multipotent hematopoietic progenitors. Functional support for niche-supporting activity was assessed using STAT5-deficient MSCs. With Lepr-Cre, niche factor mRNAs were downregulated by STAT5ab deletion with validation of reduced IGF-1 and CXCL12 proteins. Furthermore, computational analyses (differential expression/co-expression) revealed a key role for STAT5ab/Cish balance with Cish strongly co-expressed in MSCs and HSCs primed for differentiation. Therefore STAT5ab-associated gene regulation supports the bone marrow microenvironment.

## Introduction

The bone marrow (BM) microenvironment is comprised of niche cells that produce some cytokines in highly localized quantities requiring proximity for cells to derive a benefit. Outside of sensing these factors, hematopoietic stem cell (HSCs) are more likely to experience long-range acting cytokines and differentiate, leave the niche, or die. Although the epigenetic program(1) within each HSC ultimately dictates cell fate, the BM anatomic niche can have a profound effect on lineage commitment and self-renewal. Understanding transcriptional regulation of HSCs and stromal cells is critical to develop new niche-based therapeutic strategies. In this study, we explored extrinsic mechanisms that support hematopoietic stem/progenitors in the BM microenvironment and identified important functions in heterotypic bone marrow cells.

Hematopoiesis is a continuum of progressive differentiation downstream of self-renewing hematopoietic stem cells. HSCs are initially platelet-primed(2, 3) and retain multipotency. With commitment, multipotent progenitors (MPPs) progressively lose CD150 (Slamf1) marker expression and myeloid potential as they progressively acquire lymphoid potential(4–6). The regulation of this process in the BM is complex and the role of the niche in support of MPPs has not been defined, although much progress in long-term repopulating (LT)-HSC support has been gained in recent years. There is compelling data that stromal cell-derived factor-1 (SDF-1; Cxcl12) and stem cell factor (SCF; Kitl) producing cells form micro-niches with potentially specialized functions in hematopoietic stem/progenitor (HSC/HPC) support(7–10). Key outstanding questions are how niche factors ensure normal hematopoiesis, whether niche factor source determines responses, and how these factors are regulated by a variety of stromal cells. Single-cell RNA sequencing studies are beginning to address heterogeneity in wild-type mice on both the HSC(11) and stromal sides(12, 13). Additional hematopoietic support roles for T-cells(14, 15), macrophages(16) (Mac), and megakaryocytes (Mks)(17) are described, but much remains to be learned about regulatory mechanisms controlling niche factors. Other than lymphoid-primed multipotent progenitor (LMPP; MPP4) responses to IL-7(18), little is known about MPP2-3 beyond an important role for circadian-regulated Tnfα(19, 20).

Signal transducer and activator of transcription-5 (STAT5) has been a focus in our lab for two decades. Our prior work with STAT5 conditional knockout using Mx1-Cre showed defects in multi-lineage differentiation and survival but also in steady-state HSC homeostasis(21). However, Mx1-Cre can also delete STAT5 in osteolineage progenitor cells resulting in reduced bone mass(22). We have more recently used Vav1(23)-Cre(24) knockout mice to address adult hematopoiesis in the absence of an interferon response required for Mx1-Cre(25). The Vav1-Cre mouse provides a tool for assessing effects on HSCs caused by STAT5 loss but it also has potential stromal expression(26). Indeed, here we report deletion in CD45^neg^ cells. Since these tools can be promiscuous to some degree in non-hematopoietic cells, we also deleted STAT5 using a suite of Cre transgenic strains and discovered a novel extrinsic role in the transient support of multipotent progenitors (MPPs). Bone marrow hematopoietic stem cell niche single cell studies have focused on gene regulation of mesenchymal stem and progenitor cell (MSPC/MSC) and HSC differentiation alone but this study combines knockout of STAT5ab within both HSC and MSC compartments for a more holistic view.

Here, single cell transcriptomic analysis was combined with STAT5ab KO mice to explore the mechanisms of STAT5ab function in hematopoiesis. Both hematopoietic and stromal compartments were analyzed to get a full picture of the bone marrow niche. Since KLS cells include both HSC and MPPs the impact of STAT5 deletion was assessed from the LT-HSC through various multipotent progenitor cells. Technology advances to deconvolute HSC/MSC from the same scRNAseq run uncovered promiscuous expression of Nes and deletion by Vav1 in KO mice. Further Cre-mediated deletion in stromal cells and validation in isolated MSCs confirmed novel hematopoiesis-supporting functions for STAT5ab by modulating differentiation priming in both hematopoietic and non-hematopoietic progenitors.

## Results

### STAT5-deficient KLS cells have aberrant myeloid lineage priming based on GSEA and trajectory analysis

To examine STAT5ab function in cells derived from Vav1+ hematopoietic cells, STAT5 deletion with Vav1-Cre was achieved and KLS cells were sorted and analyzed by single cell RNA seq. Sequencing data from 2841 WT cells and 7167 KO cells were obtained (KLS-WT, average of total UMIs = 10979; KLS-WT, average number of genes = 3336; KLS-KO, average of total UMIs = 14715; KLS-KO, average number of genes = 3784). The subsequent data analyses yielded 13 clusters (12 hematopoietic, 1 stroma (Pdgfra^+^, Lepr^+^)) which is equivalent to 94% of purity for hematopoietic cells (**Fig. S1A**) and deletion of STAT5ab was observed throughout all the hematopoietic clusters (**Fig. S1B**) except for cluster 4 which on closer examination was revealed to be CD45^neg^. However, STAT5ab expression was mostly negative in cluster 4 so it was not possible to adequately compare Vav1-Cre/+ wild-type control cells with STAT5ab knockout cells. However, interestingly the few expressing cells had higher levels than in the hematopoietic clusters. Notably, cluster 2 was unique to STAT5ab KO whereas cluster 3 was unique to WT cells.

**Fig. 1** shows the results of UMAP (**Fig. 1A**) and trajectory analysis (**Fig. 1B**) as well a heat map of cluster-defining genes (**Fig. 1C**) with selected genes shown.

**Fig. 1.**
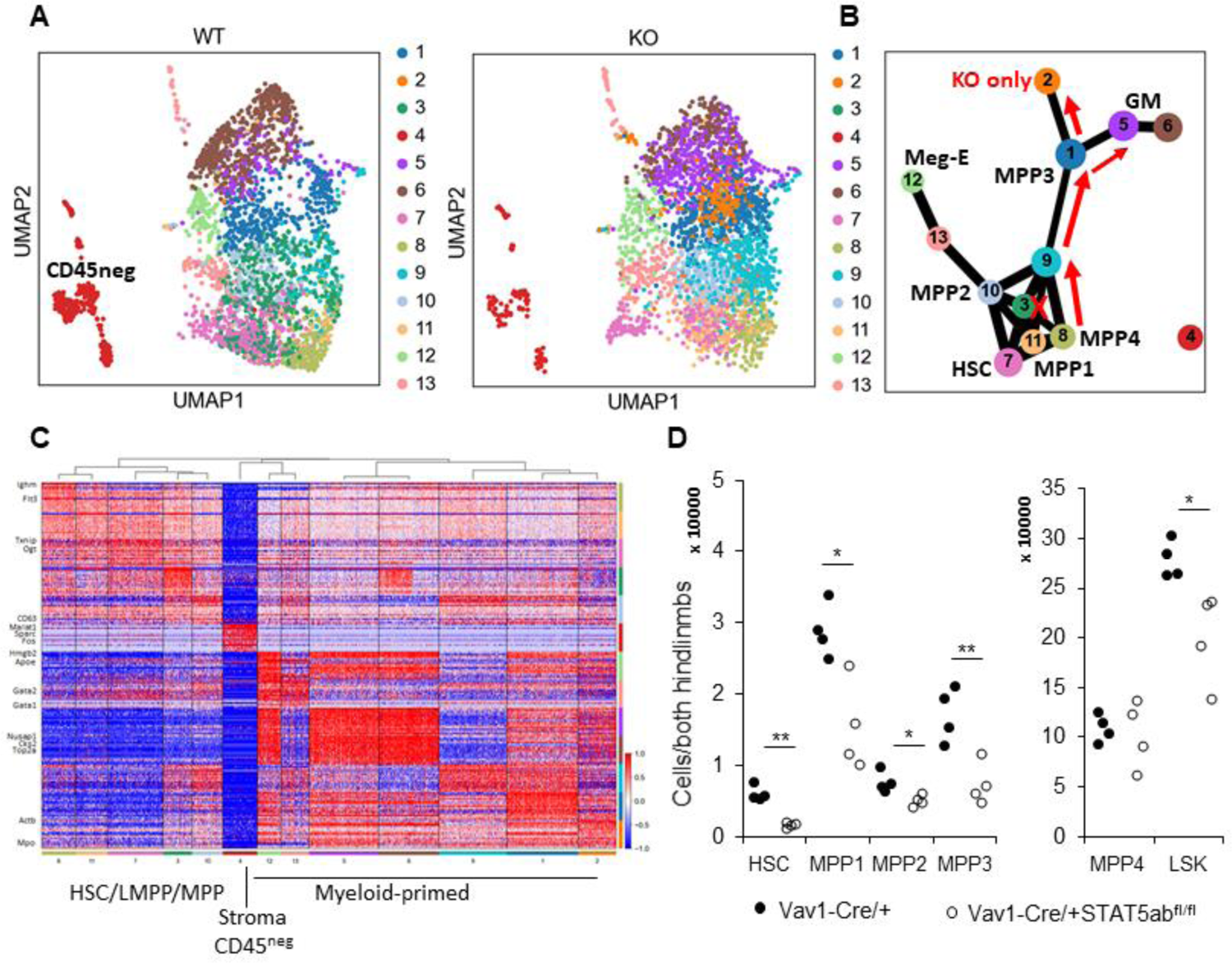
Single cell RNA seq analysis reveals STAT5-deficient KLS cells are primed for myeloid lineage differentiation. (**A**) UMAP for thirteen clusters from WT and STAT5 KO KLS cells. (**B**) Trajectory analysis of KLS single cell data. (**C**) Cluster of signature genes. Expression of the top differentially expression genes (rows) across the cells (columns) in each cluster (color bar, bottom and right, as in A and B). Key genes highlighted on left. (**D**) Conditional deletion of STAT5ab with Vav1-Cre significantly altered the population of stem and progenitors in the bone marrow cells. Bone marrow cells from Vav1-Cre/+STAT5ab^fl/fl^ and Vav1-Cre/+ control were assayed by multi-parameter flow cytometry to quantitate the number of stem and progenitors. BM cells were stained with antibodies against lineage markers, c-Kit, Sca-1, CD150, CD48, CD135, CD34, and IL7R. The absolute number of stem and progenitor cells in Vav1-Cre/+STAT5ab^fl/fl^ and Vav1-Cre/+ control mice bone marrow samples is shown (n=4 for both groups, *, p≤0.05: **, p≤0.01).

**Fig. S2** shows some key cluster-defining genes within each cluster.

Interestingly, a unique off-ramp signature was observed in cluster 2 of knockout mice where the gene signature is suggestive of osteoclasts. Bone marrow multipotent progenitors were assessed and it was found that Vav1-Cre/+STAT5ab^flox/flox^ mice had significantly reduced numbers of KLS, the most immature subsets, MPP1, and myeloid biased subsets MPP2 and MPP3 but not lymphoid-primed MPP4 cells (**Fig. 1D**).

GSEA using top differentially expressed genes from KLS cells of cluster 7 (HSC cluster) showed increases in G2-M checkpoint and reactive oxygen species (ROS) pathway genes, along with massive increases in cytoplasmic protein translation (**Table S1**).

In contrast, oxidative phosphorylation and glycolysis gene signatures were decreased. Because translation is one of the most energy consuming processes in the cell this is an unfavorable ratio for normal myelopoiesis (**Tables S2, S3**).

Increased translation but decreased metabolism is a scenario for inefficient myelopoiesis. Indeed, despite myeloid priming, pathway analysis showed priming for endoplasmic reticulum (ER) stress/unfolded protein response (UPR).

### Vav1-Cre is promiscuous with STAT5ab deletion in bone marrow CD45^neg^ stromal cells and altered differentiation and niche factor support gene signatures

To correlate expression levels and Cre-mediated deletion efficiency in bone marrow hematopoietic and stromal (Cluster 4) cell types the data was analyzed further. Vav1 was expressed in a small minority of cluster 4 cells (potential stromal cell contaminants), including in 108 Pdgrfa^+^ cells which were exclusively stromal (**Fig. 2A**). STAT5ab deletion was observed in Vav1^+^, Nes^+^, Pdgfra^+^, and Ly6a^+^ cells (**Fig. 2B**) but not in Lepr^+^ or Acta2^+^ cells of cluster 4 (**Fig. 2C**).

**Fig. 2.**
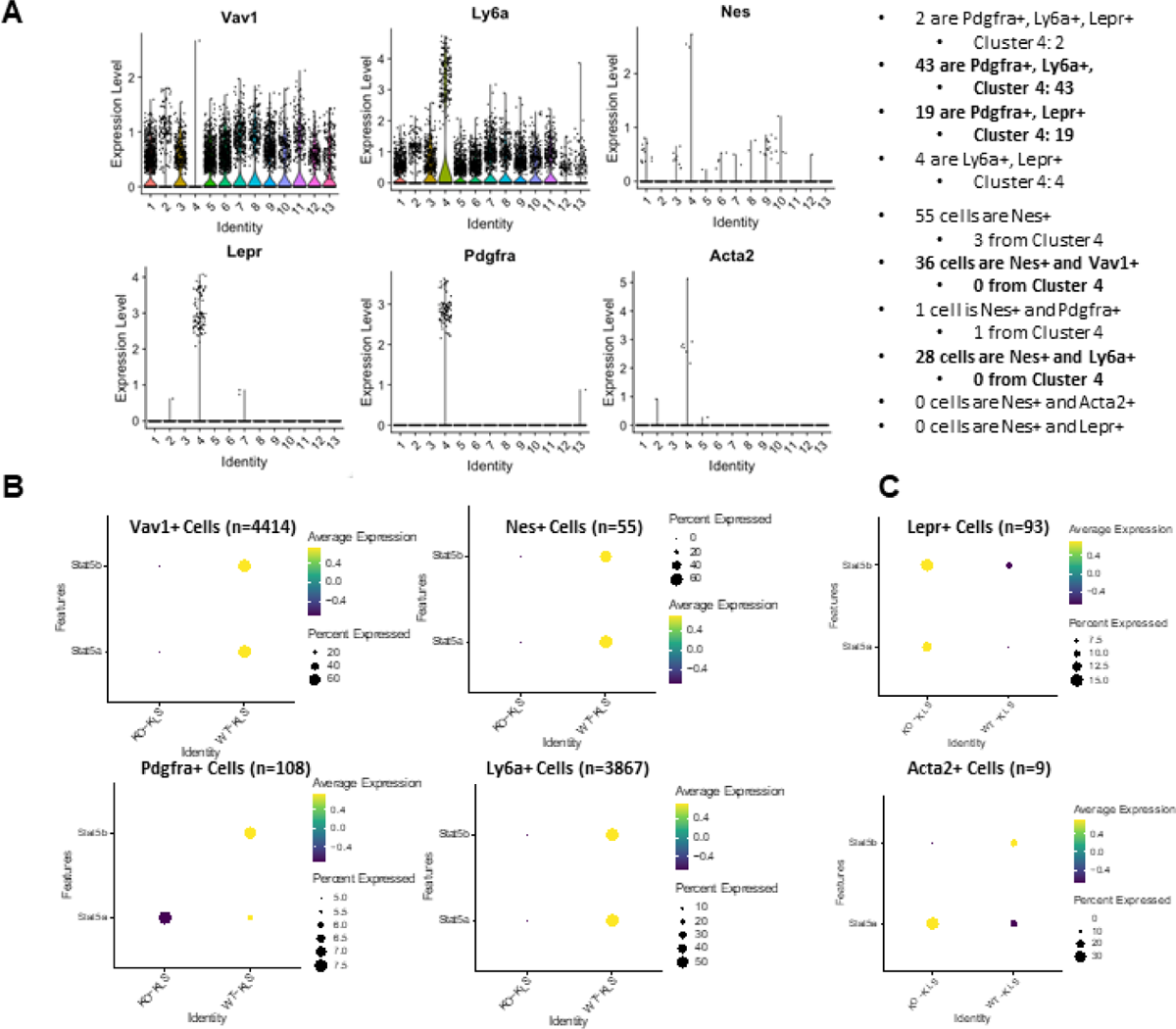
Vav1 is promiscuous in bone marrow CD45^neg^ stromal cells and Nes is promiscuous in bone marrow HSCs. (**A**) Violin plot shows the expression of distribution of Vav1, Ly6a, Nes, Lepr, Pdgfa and Acta2 across thirteen clusters. (**B**) Stat5a and Stat5b deletion efficiency is shown for Vav1-Cre mediated deletion in sorted KLS cells. The average expression and the percentage of Stat5a and Stat5b expressed between KO and WT KLS in Vav1^+^, Ly6a^+^, Nes^+^ and Pdgfra^+^ cells is shown. (**C**) The average expression and the percentage expressed of Stat5a and Stat5b between KO and WT KLS in Lepr^+^ or Acta2^+^ cells in KO KLS compared to WT KLS cells is shown.

These data show that Nes and Vav1 are co-expressed in a small percentage of hematopoietic cells and that STAT5ab was efficiently deleted in cells expressing Nes. This result suggests that Nes-Cre can delete STAT5ab in hematopoietic cells. In contrast, Vav1-Cre may be expressed more during stromal cell development leading to efficient deletion in Pdgfra^+^ cells, despite not being highly expressed in adult stromal cell types.

Given the properties of Vav1-Cre mediated deletion in KLS cluster 4, possible stromal cell contribution to the myeloid skewed hematopoietic phenotype was examined by GSEA on this cluster. Niche factor genes were decreased (Cxcl12, Kitl, Igf-1, Ntn1) suggesting that cluster 4 is comprised of CXC chemokine ligand (CXCL)12-abundant reticular (CAR) cells which are defined by high levels of Cxcl12/Sdf1 expression. Vav1-Cre cluster 4 also showed additional gene changes associated with decreased TGFβ response, bone morphogenetic protein (BMP) signaling, Wnt signaling, blood vessel morphogenesis but increased expression of negative regulators of TGFβ pathway, cartilage and bone development, and negative regulation of vascular development. Skewed toward bone/cartilage gene priming was also observed.

### Stromal Cre mouse strains reveal Nes is promiscuous in HSCs but STAT5ab deletion in MSCs decreases hematopoietic progenitor numbers *in vivo*

We previously showed that STAT5ab deletion reduced lymphocytes and resulted in a mild anemia. Now using a series of new Cre transgenic mice we tested more direct effects of STAT5ab deletion in MSCs by comparing a variety of Cre transgenic knockouts to the Vav1-Cre knockout control. **Table 1** shows the results of peripheral blood hematology analysis from Nes and Lepr-cre, transgenic mice as drivers to knockout STAT5ab. The other transgenic mice Prrx1, Osx1, and Ocn-Cre used to delete STAT5ab are listed in **Table S4**.

**Table 1.** Peripheral blood hematology of Vav1-Cre/+, Vav1-Cre/+STAT5ab^fl/fl^, Nes-Cre/+, Nes-Cre/+STAT5ab^fl/fl^, Lepr-Cre/+, Lepr-Cre/+STAT5ab^fl/fl^ mice.

|  | Vav1-Cre/+<br>(n=11) | Vav1-Cre/+<br>STAT5ab<sup>fl/fl</sup><br>(n=7) | Nes-Cre/+<br>(n=7) | Nes-Cre/+<br>STAT5ab<sup>fl/fl</sup><br>(n=12) | Lepr-Cre/+<br>(n=9) | Lepr-Cre/+<br>STAT5ab<sup>fl/fl</sup><br>(n=17) |
| --- | --- | --- | --- | --- | --- | --- |
| WBC (10<sup>3</sup>/μl) | 11.0 ± 2.1 | 1.5 ± 0.6** | 9.5 ± 1.8 | 8.2 ± 2.3 | 8.4 ± 1.5 | 9.1 ± 1.9 |
| LYM (10<sup>3</sup>/μl) | 9.4 ± 1.7 | 0.7 ± 0.5*** | 8.3 ± 1.5 | 6.5 ± 1.6* | 7.1 ± 1.5 | 7.4 ± 1.8 |
| MONO (10<sup>3</sup>/μl) | 0.5 ± 0.1 | 0.3 ± 0.0** | 0.4 ± 0.1 | 0.4 ± 0.2 | 0.5 ± 0.1 | 0.5 ± 0.1 |
| GRAN (10<sup>3</sup>/μl) | 1.1 ± 0.3 | 0.4 ± 0.1** | 0.9 ± 0.2 | 1.3 ± 0.7 | 0.8 ± 0.2 | 1.2 ± 0.5* |
| LYM % | 85.5 ± 1.9 | 51.0 ± 9.8*** | 86.7 ± 1.3 | 80.2 ± 5.2** | 84.2 ± 4.3 | 81.1 ± 5.6 |
| MONO % | 4.2 ± 0.2 | 13.6 ± 2.5*** | 4.0 ± 0.6 | 4.7 ± 0.9 | 4.9 ± 1.3 | 4.9 ± 0.7 |
| GRAN % | 10.3 ± 1.8 | 35.4 ± 8.6*** | 9.3 ± 1.1 | 15.1 ± 4.4** | 10.8 ± 3.1 | 14.0 ± 5.0 |
| HCT (%) | 44.5 ± 1.4 | 33.8 ± 4.9*** | 45.2 ± 1.1 | 45.1 ± 1.5 | 45.0 ± 2.8 | 44.6 ± 1.8 |
| MCV (fl) | 44.0 ± 0.6 | 44.2 ± 2.8 | 43.2 ± 0.5 | 44.1 ± 1.0* | 44.7 ± 1.3 | 45.4 ± 0.8 |
| RDW% | 17.0 ± 0.4 | 19.3 ± 1.5** | 17.6 ± 0.5 | 17.0 ± 0.5 | 17.2 ± 0.6 | 17.2 ± 0.5 |
| HGB (g/dl) | 15.1 ± 0.4 | 11.6 ± 1.4*** | 15.7 ± 0.4 | 15.2 ± 0.5 | 15.4 ± 0.8 | 15.3 ± 0.6 |
| MCHC (g/dl) | 34.0 ± 0.3 | 34.5 ± 0.8 | 34.8 ± 0.2 | 33.8 ± 0.5*** | 34.3 ± 0.7 | 34.3 ± 0.3 |
| MCH (pg) | 15.0 ± 0.2 | 15.2 ± 0.6 | 15.0 ± 0.2 | 14.9 ± 0.3 | 15.3 ± 0.5 | 15.6 ± 0.2 |
| RBC (10<sup>6</sup>/μl) | 10.1 ± 0.3 | 7.6 ± 0.7*** | 10.4 ± 0.3 | 10.2 ± 0.4 | 10.1 ± 0.7 | 9.8 ± 0.4 |
| PLT(10<sup>3</sup>/μl) | 299.9 ± 75.2 | 305.1 ± 62.9 | 249.1 ± 46.6 | 281.2 ± 73.1 | 289.4 ± 88 | 315.4 ± 66.4 |
| MPV (fl) | 5.7 ± 0.3 | 5.6 ± 0.2 | 5.4 ± 0.4 | 5.5 ± 0.3 | 5.3 ± 0.1 | 5.3 ± 0.2 |

\*P ≤ 0.05; \*\*P ≤ 0.01; \*\*\*P ≤ 0.001; unpaired t-test relative to each cre control group. Numbers of mice are indicated in the table. WBC, white blood cells; LYM, lymphocytes; MONO, monocytes; GRAN, granulocytes; HCT, hematocrit; MCV, mean corpuscular volume; RDW, Red cell distribution width; HGB, hemoglobin; MCHC, mean corpuscular hemoglobin concentration; MCH, mean corpuscular hemoglobin; RBC, red blood cells.

There were some notable changes in lymphocyte, granulocyte, and red blood cell indices. These changes are visible but much milder when STAT5ab was deleted by Nes-Cre compared to Stat5ab deletion by Vav1-Cre. However, the trend of those changes included a reduced percentage of lymphocytes and an increased percentage of granulocytes when STAT5ab were knockout by Lepr-Cre and Prrx1-Cre. Overall, the peripheral blood myeloid hematology was relatively normal indicating that counterbalanced myelopoiesis-promoting cytokine signaling may be active.

To assess multi-lineage repopulating activity, functional competitive repopulation assays were performed for Vav1 control, Nes, and Lepr STAT5ab knockout mice (**Fig. 3A/B**). Only Nes-Cre resulted in significantly reduced donor %CD45.2 (Fig. 3A) as well as B220^+^, Ter119^+^, and CD4^+^ cells in peripheral blood (**Fig. 3B**). Myeloid lineage hematopoiesis was rescued in these mice presumably due to a myeloid expansion specific to that lineage. A mild increase in multi-lineage competitive repopulation was observed with Lepr-Cre mediated STAT5ab deletion. In the bone marrow compartment, KLS cells were examined by flow cytometry and only Nes-Cre resulted in a significant reduction of donor contribution (**Fig. 3C**). Bone marrow competitive activity assay was also performed using STAT5ab knockout mice driven by Prrx1, Osx1 **(Fig S3A/B)**, and Ocn-Cre **(Fig S3C/D)**. All those mice had similar bone marrow competitive activity, although Prrx1 knockout had a subtle reduction in overall %CD45.2 with no significant changes at the level of multi-lineage engraftment.

**Fig. 3.**
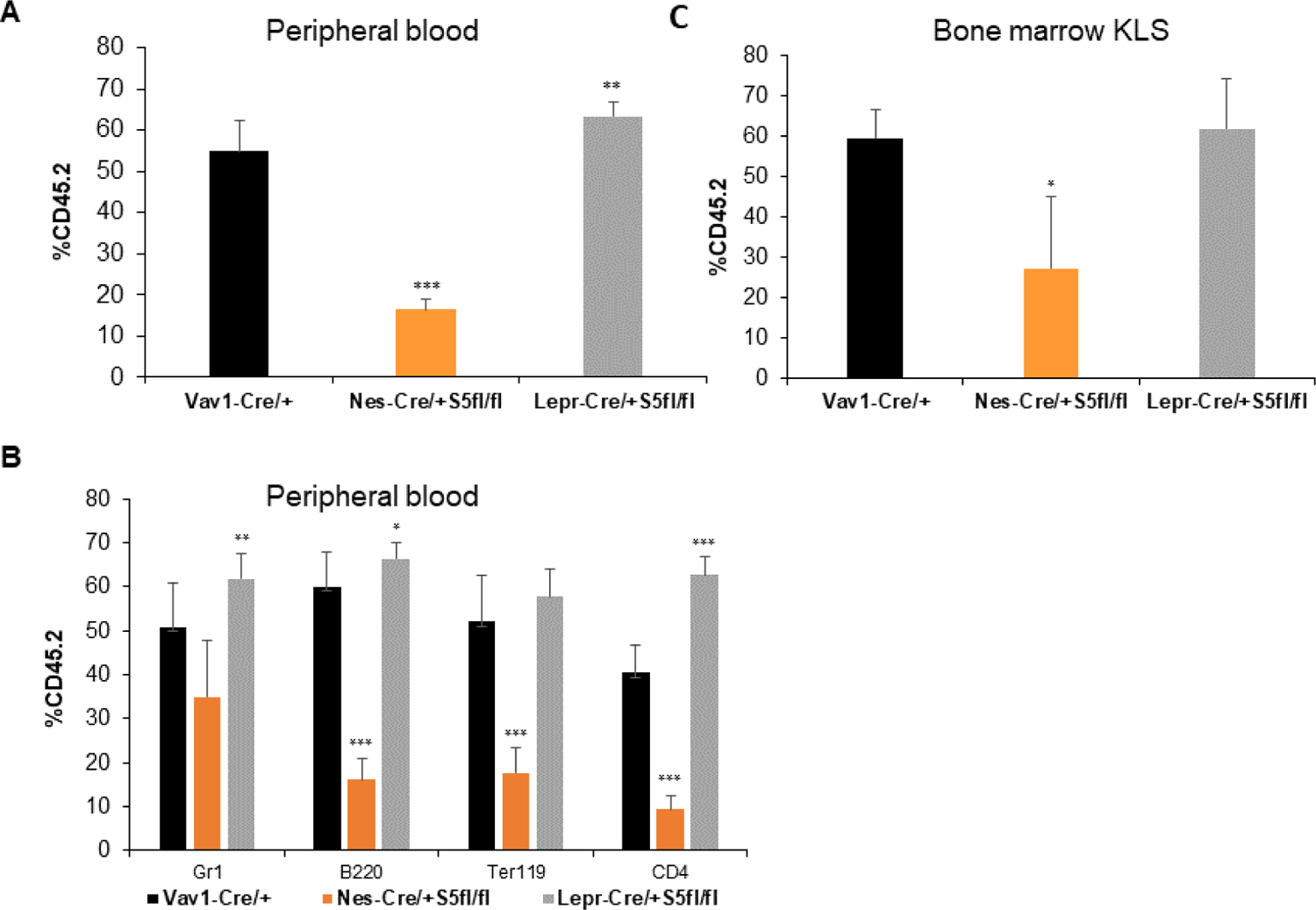
Conditional Nes-Cre deletion of STAT5ab leads to decline in mouse HSC activity while Lepr-Cre deletion has a reciprocal effect. Bone marrow cells from 3-4 donor mice were harvested from each group and mixed 1:1 with wild type Boy J competitor (CD45.1) and transplanted into lethally-irradiated Boy J mice (CD45.1) for the competitive repopulation assay. Recipient mice were bled 16 weeks later for flow cytometry analysis. Shown is the average of two independent experiments with 5 mice per group in each experiment. The genotype STAT5ab^fl/fl^ is also represented here as S5fl/fl because of space. (A) Overall donor percentage of CD45.2 positive in the peripheral blood by FACS analysis. (B) Multi-lineage analysis for the same two independent experiments. (C) Percentage of donor engraftment in the KLS fraction in the bone marrow cells.

To understand the role of STAT5ab in multipotent progenitors, mouse bone marrow cells was were analyzed by multi-parameter flow cytometry analysis. KLS cells were subdivided into four populations based on the expression of CD150 and CD48: CD150^+^CD48^neg^KLS (HSCs) (**Fig. 4SA**), CD150^neg^CD48^neg^KLS (MPPs) (**Fig 4SB**), CD150^neg^CD48^+^KLS (HPC-1) (**Fig. 4SC**), and CD150^+^CD48^+^KLS (HPC-2) (**Fig. S4D**).

Multipotent progenitors are known to be heterogeneous. The Trumpp(28, 29) and Passegue(4) groups divided the MPP population into MPP1-4 according their immunophenotype using CD34, CD135, CD150, CD48 along with KLS markers. Using those markers, Lepr-Cre and Nes-Cre STAT5ab knockout mice were examined more thoroughly to better understand the effects of Cre mediated deletion of STAT5ab on the population of stem and progenitors. Nes-Cre/+STAT5ab had consistent reduction of long term HSCs (CD34^neg^CD48^neg^CD150^+^CD135^neg^KLS) (**Fig. 4A**) while STAT5ab deletion with Lepr-Cre was associated with accumulation of HSCs at the expense of MPP4/CLP lineage (**Fig. 4B**) as determined by flow cytometry. This result is consistent with a recent publication showing how competition between hematopoietic stem and progenitor cells controls HSC compartment size(30). Therefore, Lepr^+^ cells utilize STAT5ab in a manner that promotes lymphoid lineage priming.

**Fig. 4.**
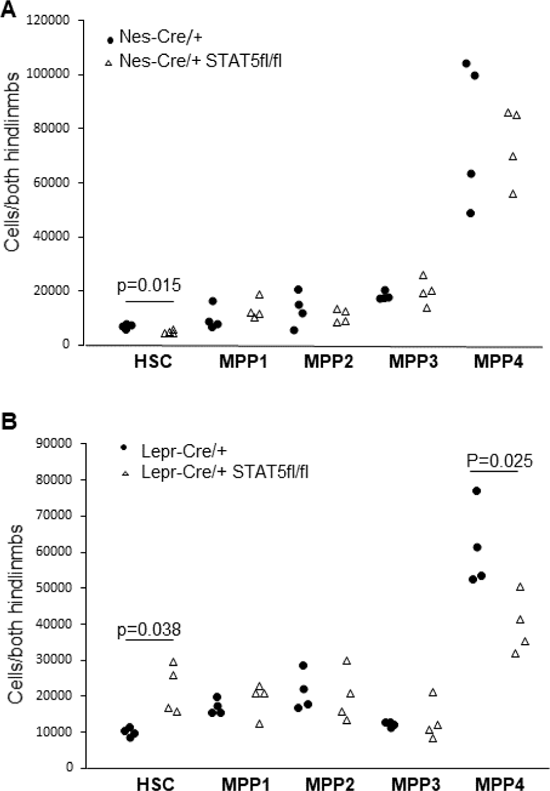
Conditional deletion of STAT5ab with Nes-Cre and Lepr-Cre has reciprocal effects on the number of HSCs in the bone marrow. Bone marrow cells from Nes-Cre/+STAT5ab^fl/fl^ and Nes-Cre/+ or Lepr-Cre/+STAT5ab^fl/fl^ and Lepr-Cre/+ control were assayed by multi-parameter flow cytometry to quantitate the number of stem and progenitors. BM cells were stained with antibodies against lineage markers, c-Kit, Sca-1(Ly6a), CD150, CD48, CD135 (FLK2), CD34, and IL7R. The genotype STAT5ab^fl/fl^ is also represented here as STAT5^fl/fl^ because of space. (**A**) Absolute number of stem and progenitor cells in Nes-Cre/STAT5ab^fl/fl^ and Nes-Cre/+ control. (**B**) Absolute number of stem and progenitor cells in LepR-Cre/STAT5ab^fl/fl^ and Lepr-Cre/+ control (n=4 for each group).

### Single cell RNA seq of CD45^neg^ stromal cells using Lepr-Cre deletion of STAT5ab reveals overlap between several MSC markers

Having defined the impact of STAT5ab deletion on the numbers of hematopoietic stem/progenitors, we next explored molecular mechanisms that might be responsible. To examine STAT5ab function in cells derived from Lepr^+^ stromal cells, STAT5ab deletion with Lepr-Cre was used and CD45^neg^Ter119^neg/low^CD71^neg^Sca-1^+^CD140a^+^ living cells were sorted and analyzed by single cell RNA seq. Sequencing data from 565 WT and 349 KO cells were obtained (MSC-WT, average of total UMIs = 2789; MSC-WT, average number of genes = 1131; MSC-KO, average of total UMIs = 2990; MSC-KO, average number of genes = 1100). The subsequent data analyses yielded 6 clusters (1 early erythroid with some lymphoid genes as well, 5 stromal) (**Fig. S7/S8**).

This result is consistent with a report that bone marrow CD45^neg^ contains erythroid and lymphoid progenitors(31). The yield of captured single MSCs was lower than obtained for the KLS during the process of single cell RNA seq and thus resulted in fewer clusters. Notably, of the 6 clusters identified, deletion of STAT5ab was observed mainly when looking at the Lepr^+^ cells but not by looking at the cluster designations despite cluster 4 being the most enriched for Lepr^+^ cells.

Clusters were annotated using a panel of partially overlapping markers including Lepr, Pdgfra, Nes, Acta2, Sca1 (Ly6a), and Bglap. The UMAP (**Fig. 5A**) and lineage trajectories (**Fig. 5B**) are shown. Notably, Pdgfra was expressed in clusters 2, 3, 4, like that of Lepr (**Fig. 5C**). Interestingly, Lepr^+^ cells mostly co-expressed STAT5b but not STAT5a and Lepr-Cre caused some deletion of STAT5b mRNA levels (**Fig. 5D**). In contrast, Nes was expressed rarely (only ∼1% of MSC sequenced) in adult stroma and appeared to mark a potential transient “activated” cell intermediate based on trajectory analysis and cluster 5. Ly6a was also expressed in these 3 clusters with an expression pattern that was reciprocal to Lepr. Vav1 was expressed in cluster 5 but not enough to explain deletion resulting in adult mice. This suggests that Vav1 may be expressed in a transient early pre-MSC that results in the deletion observed in the KLS cluster 4. Lepr-Cre deleted very well in defined Lepr^+^ cells, although these cells were a minority of the cluster that we defined as the Lepr^+^ cluster because of the highest Lepr expression levels. Notably, Lepr-Cre also deleted in Pdgfra^+^ cells but not in Nes^+^, Ly6a^+^, or Acta2^+^ cells (**Fig. 5D**).

**Fig. 5.**
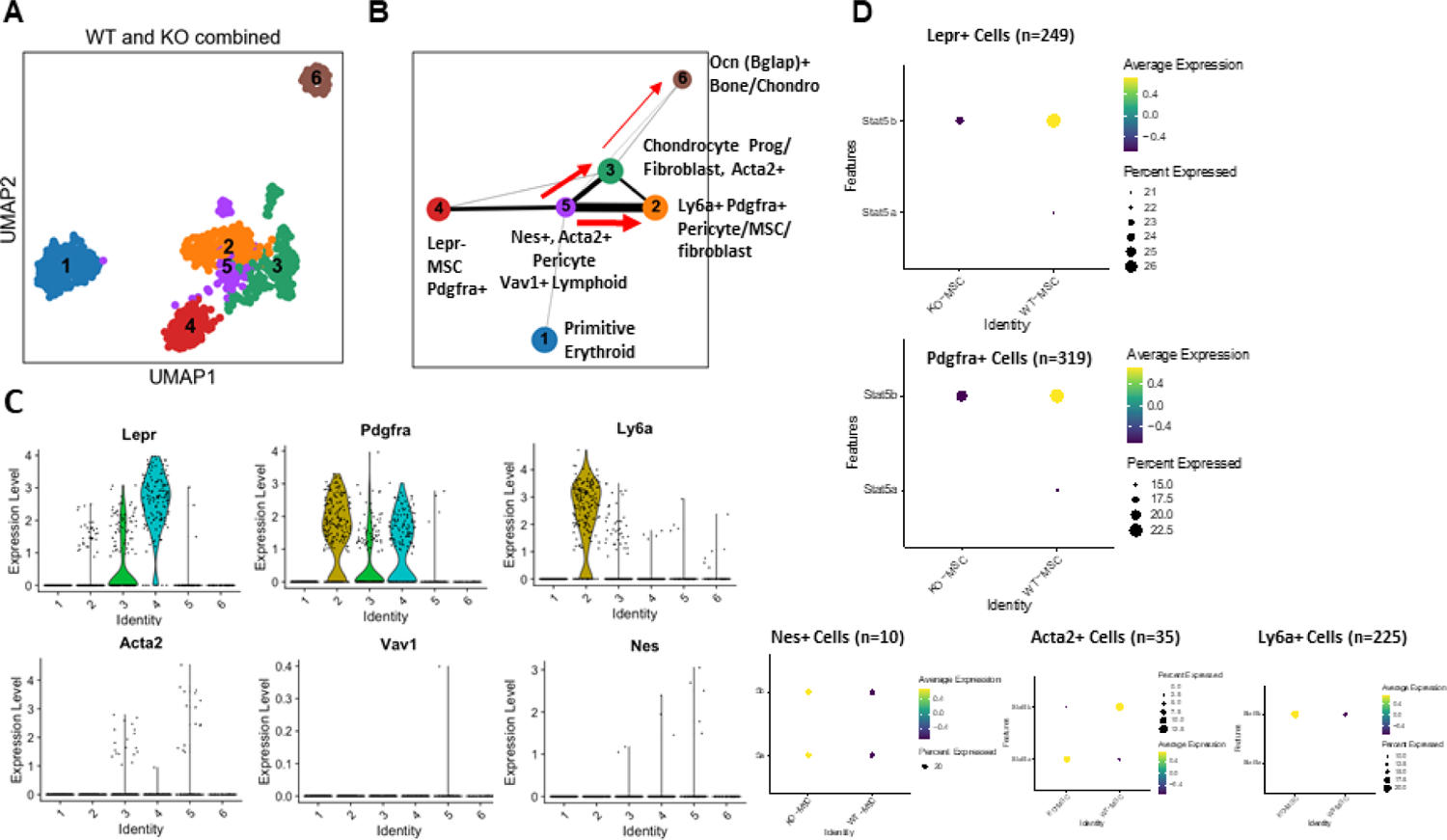
Single cell RNA seq of CD45^neg^ stromal cells using Lepr-Cre deletion of STAT5ab reveals overlap between several MSC markers Single cell RNA seq analysis was performed on stromal cells from Lepr-Cre/+Stat5ab^fl/fl^ and its control. (**A**) UMAP for six clusters of stromal cells single cell RNA seq data. (**B**) Trajectory analysis of single stromal cells by RNA seq data. (**C**) Distribution of expression level for selected marker genes across six clusters with violin plot representation. (**D**) Stat5a and Stat5b average expression and percent expressed among selected marker gene positive cells between Lepr-Cre/+ STAT5ab^fl/fl^ (KO-MSC) and its control (WT-MSC).

### Niche factor expression and support activity depends on STAT5ab expression

Analysis of the scRNAseq data set for Lepr-Cre mediated STAT5 deletion in stroma showed a strong signature of decreased niche factor genes such as Igf1, Cxcl12, Igfbp4, Igfbp7, Il34, Pdgfa, Vegfa (**Fig. 6A**) in cluster 4. Since decreased local production of Igf1 in the bone marrow microenvironment causes many hallmarks of LT-HSC aging, including cell cycling, this may explain some of the LT-HSC accumulation observed with Lepr-Cre. Reciprocally, Grem1 was the 6^th^ most induced gene in the MSCs which is a known marker for loss of multipotency (**Fig. 6A**). Cluster 5 wasn’t as changed as other clusters and this correlated with weaker STAT5ab expression levels in WT. It did trend toward more osteoblast differentiation genes and myeloid cell differentiation genes with loss of negative regulation of MAPK, p38 MAPK, ERK1/2 pathways. Other gene sets increased included regulators of osteoblast differentiation, BMP response, fat cell differentiation, as well as early response genes (fosb, jun, junb, fos, ebf1, and dusp1). Overall, the gene signature in cluster 4 as well as in cluster 2 suggests more priming toward osteo/chondro-genic differentiation (**Fig. 6B**) very similar to what was observed with the KLS cluster 4 (the only CD45neg cluster). In contrast, clusters 3 and 6 became very chondrocyte-primed (**Fig. 6C**). Altogether, these data indicate that STAT5ab deficiency may lead MSCs toward a terminal chondrocyte fate which is associated with different anatomic localization in the bone and movement away from the primary bone marrow niche that is supportive of multipotent progenitors (**Fig. 6D**).

**Fig. 6.**
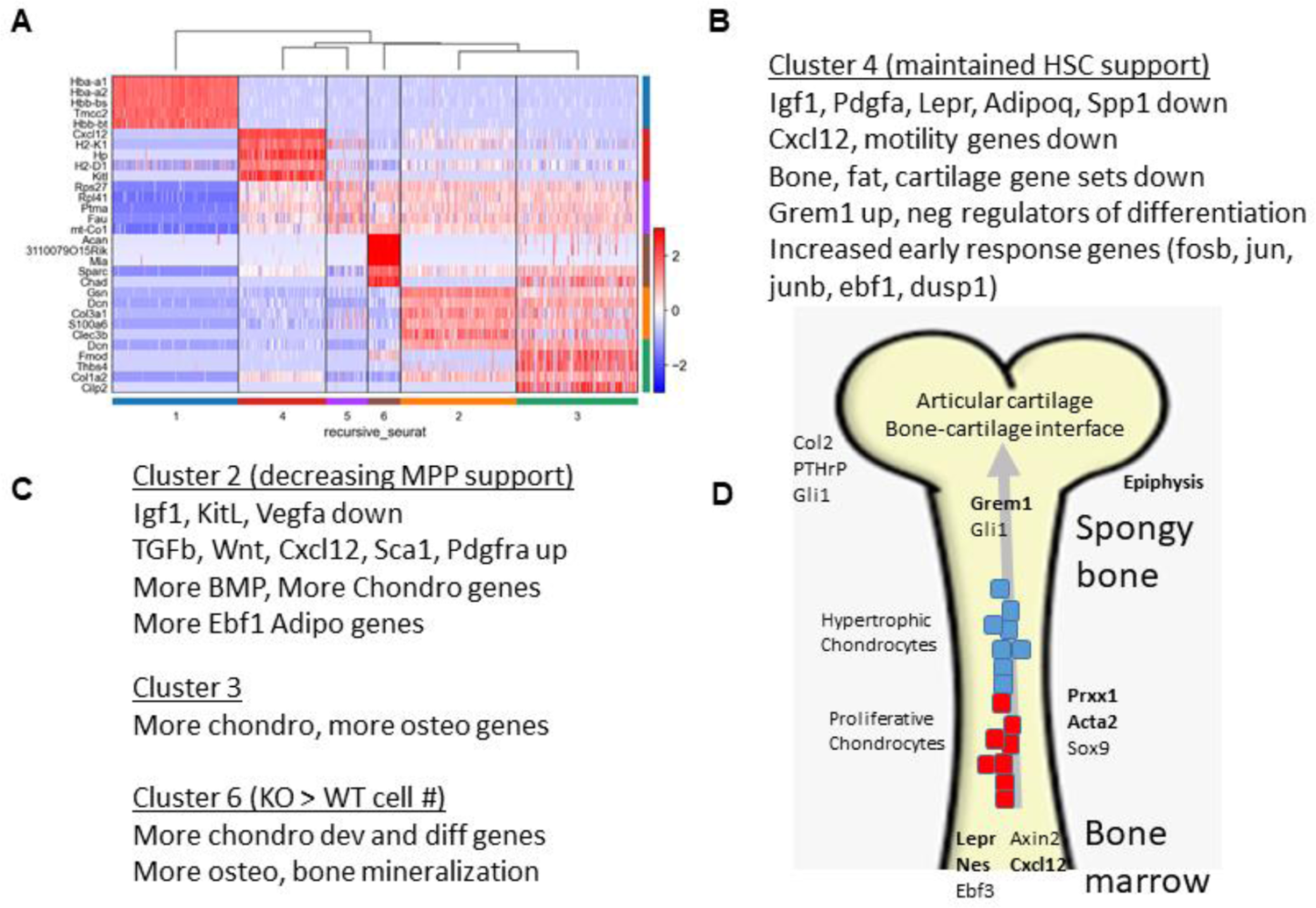
Analysis of the scRNAseq data set for Lepr-Cre mediated STAT5 deletion in stroma shows dysregulated osteo-chondro-genic differentiation and hematopoietic progenitor support functions. (A) Cluster of signature genes that define each of the MSC clusters. Expression of the top differentially expression genes (rows) across the cells (columns) in each cluster (color bar, bottom and right, as in A and B). Key genes highlighted on left. (B) The gene signature in Cluster 4 maintained HSC and MPP support. (C) The gene signature in the other stroma clusters. (D) Diagram of the molecular changes occurring during osteo-chondro-genic differentiation in the anatomic context of the bone and bone marrow space. All gene sets were identified from Panther GSEA as significantly changed and P<<0.05.

To investigate whether these changes are observed in cultured MSCs *in vitro*, MSCs were analyzed by qRT-PCR for reduced mRNA and protein for STAT5a and STAT5b separately (**Fig. 7A/B**). The key gene from the global transcriptome analysis that was increased was Grem1 and likewise it was sharply increased in cultured MSCs (**Fig. 7C**). In contrast, Cxcl12, Angpt1, Igfbp4, Igf1, and Il34 were all decreased like the global scRNAseq data (**Fig. 7D)**. Also, similarly increases in Kitl were observed. Functional analysis of STAT5ab-deficient MSCs were precluded by difficulties in culturing them following deletion for prolonged time. Therefore, to test trophic function *in vitro*, mouse embryonic fibroblasts lacking STAT5ab were used (**Fig. 7E/F**). Support function for the SCF-dependent cell line EML was tested by co-culture on STAT5ab-deficient MEFs. Reductions in support were observed compared to wild-type MEFs, consistent with a functional role for STAT5ab in providing necessary trophic support to hematopoietic progenitors. To check the production of the secretory niche factors, wild-type or STAT5ab-deficient MEFs or MSCs were cultured. The same number of cells (0.1 million) were seeded in a 24-well plate. The second day, the fresh medium with reduced FBS (2%) was changed and the supernatant was harvested 24 hr later for IGF-1 and SDF-1α ELISA assay. MEFs had a relatively lower level of basal IGF-1 and SDF-1α compared with MSCs but both STAT5-deficient MEFs (**Fig. 7G**) and MSCs **(Fig. 7H)** had significantly reduced secretory niche factors IGF-1 and SDF-1α.

**Fig. 7.**
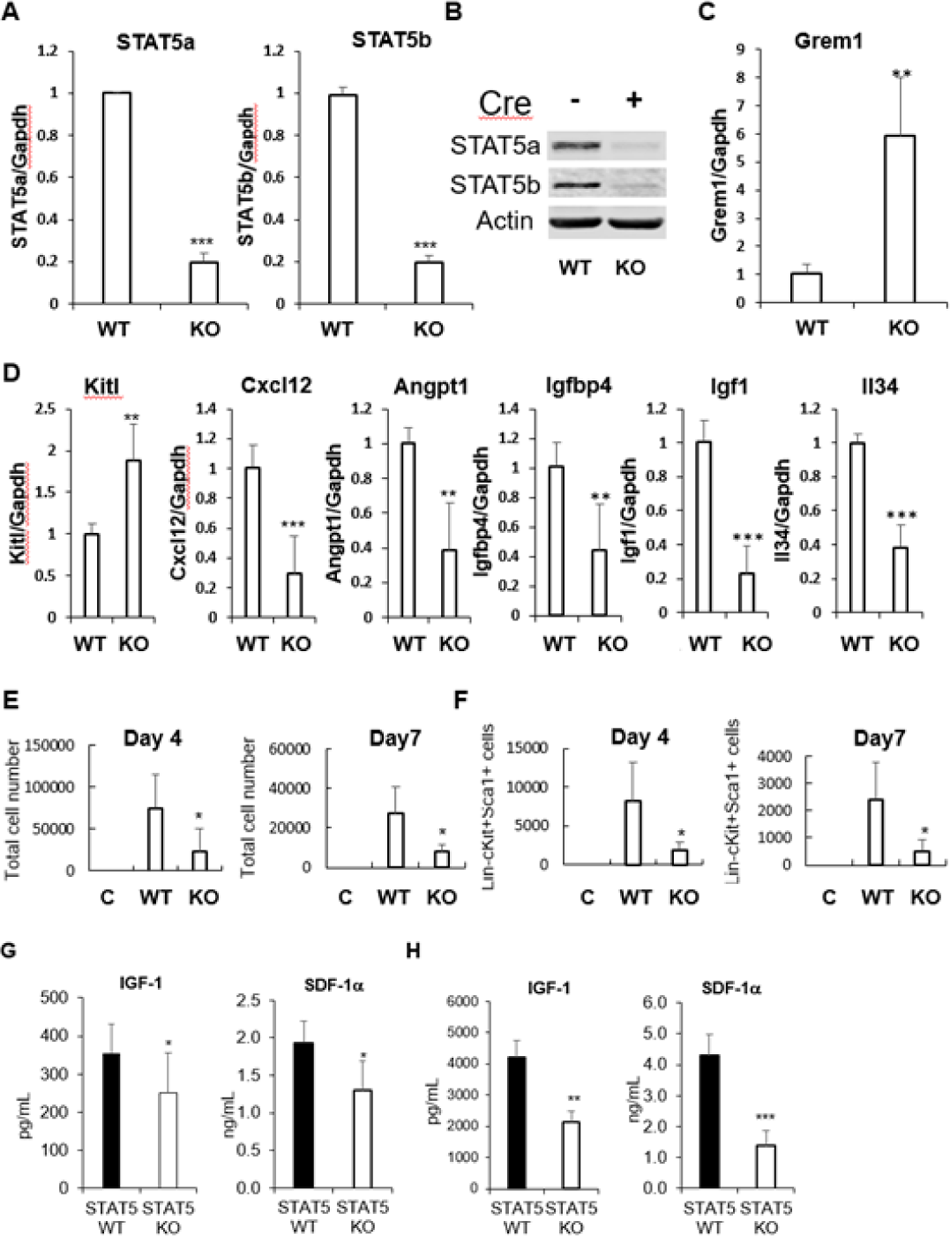
Deletion of STAT5ab with Lepr-Cre leads to alteration of bone marrow niche factor expression. **(A)** Real-time PCR showed the deletion of Stat5a and Stat5b in MSCs from Lepr-Cre/+STAT5ab^fl/fl^ compared to the MSCs from control mice. GAPDH was used as the internal control for real-time PCR. **(B)** Western blot showed the deletion of STAT5a and STAT5b protein. Actin was used for the loading control. **(C, D)** Expression of bone marrow niche factor including Grem1, KitL, Cxcl12, Angpt1, Igfbp4, Ifg1, Il34 by real-time PCR compared WT MSCs with STAT5ab deleted MSCs. **(E, F)** STAT5ab^null/null^ MEF cells have reduced hematopoietic-supporting activity for EML C1 cells. The total number of EML C1 that were co-cultured for 4 days (n=4, p=0.04) or 7 Days (n=4, p=0.03), and for Lin^neg^Kit^+^Sca-1^+^ cells co-cultured for 4 days (n=4, p=0.05) or 7 days (n=4, p=0.04) were compared to those co-cultured with WT MEF cells or without MEF (C: control). Niche factor SDF-1α and IGF-1 are significantly reduced when STAT5ab is deleted in MEF and MSCs. (**G**) Conditioned medium from wild type and STAT5ab^null/null^ MEFs was assayed for SDF-1α and IGF-1 by ELISA assay (n=5, p= 0.02 for SDF1α; n=11 p=0.01 for IGF-1). (**H**) Conditioned medium from wild type and STAT5ab deleted MSCs was assay for SDF-1α and IGF-1 by ELISA assay (n=5, p<0.001 for SDF1α, p=0.002 for IGF-1).

**Fig. 8.**
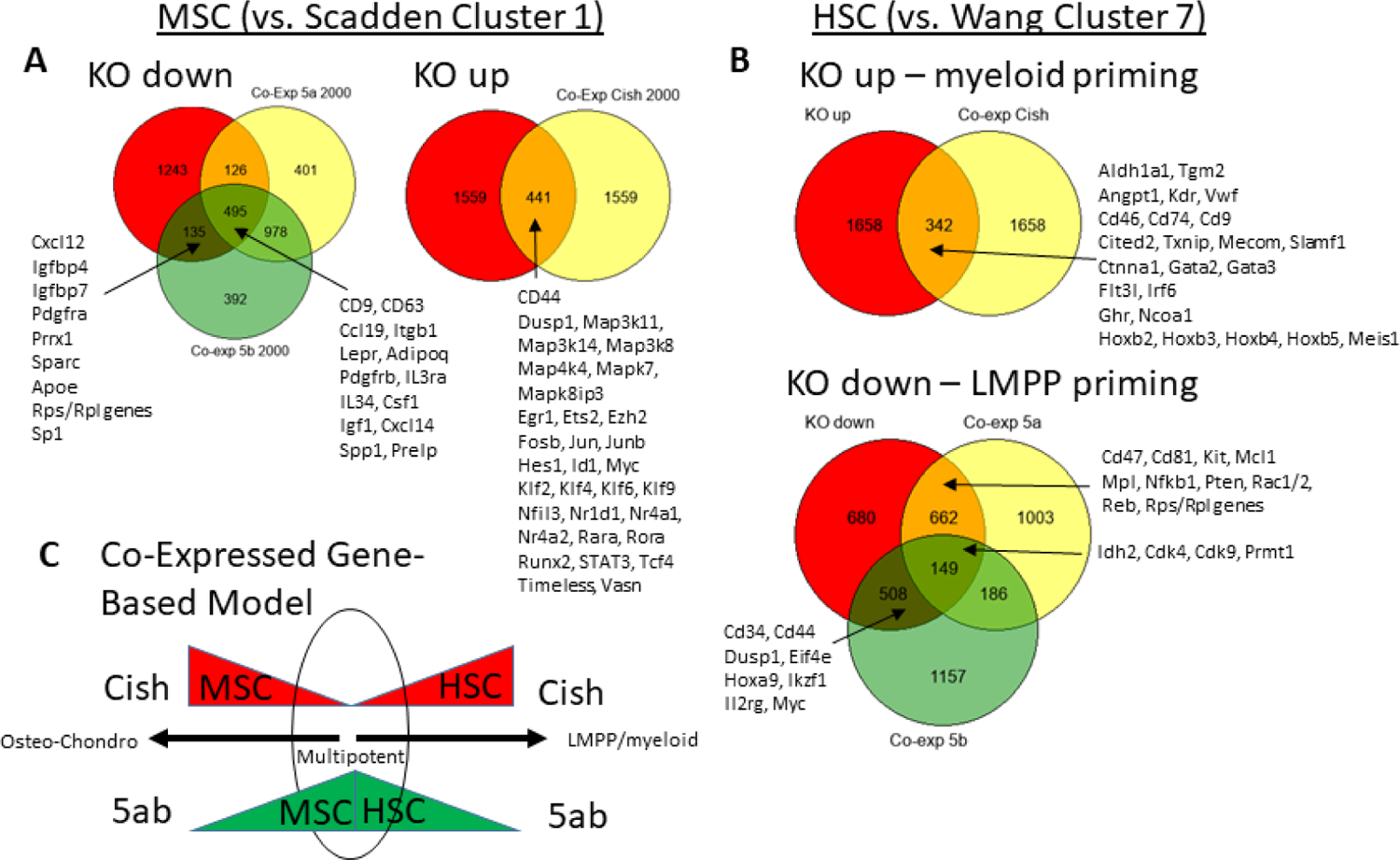
Combined STAT5ab knockout and wild-type co-expression analyses identify common STAT5ab/Cish regulation of MSC/HSC differentiation. (A). The Scadden data set was used to interrogate Lepr-MSCs compared with our wild-type and STAT5ab knockout data set. Overlap was identified by Venn Diagrams comparing the highest scoring genes in the Scadden data set with the most co-expressed genes in our data set. The most increased or decreased gene sets were analyzed separately to discover new signatures associated with gain or loss of STAT5ab expression level. (B). Similar to (A) but using our KLS data set, both knockout and wild-type gene sets were compared by Venn diagrams to determine those highest co-expressed genes that were also the most increased or decreased by Stat5ab knockout. (C). Summary of the results from the first two panels shows how STAT5ab/Cish may function in a reciprocal manner as a general biomarker of hematopoietic and mesenchymal lineage differentiation within the bone marrow.

### STAT5ab/Cish co-expression algorithms define gene signatures of differentiation priming in both MSCs and HSCs

Since all of our work had explored STAT5ab double KO mice, potential individual roles for STAT5a or STAT5b could not be separated. To attempt to address this computationally and to develop a new algorithm for co-expression analysis of STAT5a or STAT5b with genes in scRNAseq data sets, we set out to validate MSC datasets published by Scadden(12) using maximum mean discrepancy (MMD) analysis in comparison with two other co-expression algorithms (Iacono(32) and MAGIC(33)) (**Fig. S9/10**).

In combination with STAT5a or STAT5b was included Cish a known direct target and negative feedback inhibitor of STAT5ab. The combination of STAT5ab/Cish proved to be a powerful combination for identifying gene signatures in bone marrow stromal cell types associated with glucocorticoid signaling and osteogenic/chondrogenic differentiation of mesenchymal stromal cells (**Fig. S11/12**).

In hematopoietic stem/progenitor cells using scRNAseq dataset published by Gottgens(11), STAT5ab/Cish unique and overlapping signatures were associated with distinct quiescence/self-renewal vs. cell cycle activation gene signatures (**Fig. S13**).

## Discussion

Signal transducers and activators of transcription (STAT)s are a family of seven transcription factors that are activated by a diverse set of receptors and Janus kinases (JAKs). Our lab(21, 25, 34–43) and others(44–46) have shown that conditional deletion of STAT5 with Mx1-Cre caused loss of quiescence and progressive loss of CD150 expression, along with impaired lymphocyte(47) and leukemia development(48, 49). Within KLS cells, deletion of STAT5ab increased LMPPs(25) at the expense of megakaryocyte/erythroid (Mk-E) priming, much like the Thpo receptor (Mpl) knockout(50). STAT5 in signaling downstream of SDF1 or SCF has also not previously been reported and STAT5 is not tyrosine but is serine phosphorylated downstream of these important niche factors(51–53).Therefore, STAT5 is modulated by combinations of phosphorylation events which in different anatomic contexts and stromal cell populations may vary and result in different outcomes. STAT5aS780 is constitutively phosphorylated regardless of starvation, the same as reported in human CD34^+^ cells(54). Molecular investigations of the phenotypic HSC compartment have also defined a metabolically activated compartment with limited self-renewal, termed MPP1, which may represent an initial differentiation step. Interestingly, among the top cluster defining genes in this study there was a high representation of translation associated gene expression in STAT5ab knockout KLS cells. These RPS/RPL genes were ranked as high, med, or low within the top cluster-defining genes. The early clusters 7, 8, 10, 11, 3 all had more cluster-defining RPS/RPL genes whereas the more Mk/E and GM clusters didn’t have any (**Fig. S2**). This likely indicates that STAT5 deficiency causes cellular activation and abnormal myeloid lineage priming in the early MPPs. In the MSC clusters, only cluster 5 is primed with RPS/RPL gene, may represent an “activated MSC” state intermediate between quiescent Lepr-MSC and the osteo/chondro progenitors. That fits the GSEA and lineage trajectory analyses.

Intrinsic metabolic reprogramming of MSCs has been described to control their cell fate. Naïve MSCs are ROS^low^ and favor glycolysis in the hypoxic niche but switch to oxidative phosphorylation (oxphos) and become ROS^high^ with differentiation(55). STAT5ab co-expressed genes were associated with priming for translation and oxphos and STAT5 may play a role to ready MSCs for lineage commitment with Cish co-expression. In contrast to reduction of pTyr dosage which may impair terminal differentiation, STAT5 protein deletion may permit more osteolineage commitment at the expense of adipocyte priming due to loss of as yet undefined non-pTyrSTAT5 regulatory functions. pTyrSTAT5 seems likely for full lineage differentiation however. Lepr-MSC1-4 differentiation continuum genes include glucocorticoid response genes, supporting the known ability of dexamethasone to induce MSC commitment to differentiation(56). Non-genomic roles for non/low pTyrSTAT5a and to a lesser extent STAT5b in golgi have also been reported(57–59) with knock-down *in vitro* resulting in golgi fragmentation. MSCs have been extensively studied *in vitro* and *in vivo* under steady-state and stress conditions (high GC) and STAT5b transcriptionally promotes osteolineage (OLC) differentiation(60), yet understanding of STAT5-mediated hematopoiesis-support is unknown. Innervated Nes^+^ pericyte MSCs respond to the sympathetic nervous system in a circadian manner to regulate secretion of niche factors such as Cxcl12(61) and nerve inury impairs hematopoiesis(62).

Extrinsic activation of STAT5ab/Cish as a key signaling cascade can also be part of the bone marrow microenvironment to regulate MSC function. A single skeletal stem cell produces chondrocytes, osteoblasts, adipocytes and fibroblasts. STAT5 primes multipotency and KO increases differentiation potential but cannot dictate terminal differentiation. The circadian response is a major regulator of hormone(63, 64) and cytokine(65) trafficking/secretion and it is significant to better understand whether STAT5 isoforms have evolutionarily conserved roles in the BM microenvironment to support niche factor production(66, 67). Since the 1990s, roles for STAT5a (mammary gland-prolactin-milk proteins) and STAT5b (liver-growth hormone (GH)-insulin-like growth factor 1 (Igf1)) have been shown to be distinct(68) despite 95% homology. During sleep, expression of GH or prolactin precedes the glucocorticoid (GC) pulse, permitting STAT5 to cooperate with the glucocorticoid receptor (GR) in MSCs. A conserved role for STAT5 in regulating hematopoiesis through bone marrow niche factors was still surprising. Stromal support function is more subtle than the direct hematopoiesis functions previously characterized. The defects specifically and transiently resulted in impaired multipotent progenitors, thus providing novel insight into MPP niches.

STAT5 roles in MSC adipocyte(69–71) and osteoclast differentiation(22, 72) have been described to come together in regulating bone development(73). However, there has not been a previous in-depth examination of STAT5ab at the single cell level as shown in this study. Gremlin1 (Grem1) is an antagonist for bone morphogenic protein (BMP) and importantly was the main differentiation marker increased in STAT5ab knockout MSCs. Osteochondroreticular (OCR) stem cells have been recently described as a population of Nes^neg^ mesenchymal precursors that can differentiate into osteoblasts, chondrocytes, reticular marrow stromal cells, but not adipocytes *in vivo*. Grem1-Cre cells can be found concentrated within the metaphysis of long bones, but not in the perisinusoidal space where the main hematopoiesis support activity occurs. Grassinger et al.(74) described a unique approach to investigate the microanatomic location of HSPCs in long bones using combinations of micro-computed tomography (CT), histomorphometry, homing, and spatial distribution assays. At short time points after transplantation in physiologic conditions individual HSCs homed preferentially to the trabecular-rich metaphysis of long bones after transplantation. There are a number of markers found in the perivascular bone marrow niche including PDGFR*β*, PDGFR*α*, CD146, Nestin, LepR, and Cxcl12, while arterial vasculature is associated with cells expressing PDGFR*β*, PDGFR*α*, Sca1, and LepR(8, 75–77). The skeletal growth plate resting zone has a completely different set of key genes including Grem1, Col2a1, PthrP and Itgav (CD45^-^/TER119^-^/Tie2^-^/Thy^-^/6C3^-^/CD105^-^/CD200^-^/Itgav^+^). Therefore, anatomically mislocalized stromal cells may underlie the molecular changes observed with STAT5ab deficiency in this study as diagramed in Fig. 6D. Defining these changes may be a future research direction.

Vav1-Cre/+ STAT5ab^flox/flox^ mice have partially overlapping gene expression changes with the Ntn-Neo1 autonomous signaling network (data not shown). This suggests that Vav1-Cre is not only promiscuous in some mesenchymal progenitor cells but may also express Cre in arteriolar endothelial or smooth muscle cells. Ntn1 is a laminin-related secreted protein this is greatly reduced in KLS Cluster 4 (stromal cluster; gene rank 521) and this ligand binds Neo1 to maintain HSC quiescence. Ntn1 is expressed in arteriolar endothelial and smooth muscle cells to regulate quiescence and its absence increases competitive repopulation and aging phenotypes. Neo1 is one of the most consistently upregulated genes found upon HSC aging. Notably, an Acta2^+^ 24-gene co-expression “Vascular Smooth Muscle Cell Network” compared to KLS Cluster 4 (20/24 genes affected), raises the question of whether KLS (Cluster 4) are Acta2^+^ smooth muscle cells. Future studies are required to examine this area further.

In summary, this study shows for the first time that STAT5ab are active in heterotypic bone marrow stem/progenitor cells to regulate hematopoiesis. We did not examine impacts on bone, cartilage, or fat cell maturation and development in this study. Further expansion of the predictive co-expression algorithm has prognostic potential for using STAT5ab/Cish regulated biological processes as a biomarker. This could be applicable for abnormal hematopoiesis (leukemogenesis) and immunity or in the setting of cancer treatment for solid tumors with a strong anti-tumor immunity. The weakness of any scRNAseq based biomarker approach is that these are currently not amenable on a large scale beyond experimental studies. Further advances in the technology and cost reductions will be needed in order to apply this to patients in a timely manner. In the meantime, more development of the algorithms and experimental studies are necessary to refine the predictive potential and identify the most promising clinical scenarios where this approach could have an impact on decision-making.

## Materials and Methods

### Transgenic and knockout mice

All animal procedures were approved by the Emory University Institutional Animal Care and Use Committee. STAT5ab^flox/flox^ (interchangeable with STAT5ab^fl/fl^ as used in the text and figures) mice were originally obtained from Lothar Hennighausen (NIDDK, NIH) and have been bred in the Bunting lab for almost 20 years. Vav1-Cre transgenic mice were generated and used as previously described. This study used the following Cre-expressing transgenic mouse lines obtained from the Jackson Laboratory (Bar Harbor, ME) to generate new stromal knockout strains lacking expression of STAT5ab: Nes-Cre/+ (#003771), LepR-Cre/Cre (#008320), Prrx1-Cre/+ (#005584), Osx1-GFP:Cre/+ (#006361), BGLAP(OCN)-Cre/+ (#019509) and Nes-CreERT2/+ (#016261). Cdh5-CreERT2/+mice generated by Ralf Adams were generously provided by Brian Petrich (Emory). To delete STAT5 in hematopoietic or mesenchymal lineages, various Cre mice were genetic crossed with STAT5ab^flox/flox^ mice. For inducible Cre (Nes-CreERT2 or Cdh5-CreERT2/+) knockout, mice were treated with Tamoxifen (MilliporeSigma, Burlington, MA). Tamoxifen was dissolved in corn oil at a concentration of 20 mg/ml by shaking overnight at 37°C in foil paper wrapped tube to protect from light and then mice were injected with 75 mg tamoxifen/kg body weight via i.p. injection once every 24 hours for 5 consecutive days. The final analysis of those mice was carried out two weeks after the last dose of treatment.

### Peripheral blood hematology

Following puncture of the retro-orbital venous sinus or alternative facial vein, the mouse peripheral blood was collected in a heparinized capillary tube (ThermoFisher Scientific, Norcross, GA). Mouse hematology was determined using a HemaTrue Hematology Analyzer (Heska, Loveland, CO). A very small percentage of Vav1-Cre/+STAT5ab^fl/fl^ mice develop leaky lymphoid and erythroid cells. Those mice are excluded from further analysis despite being knockout by genotyping.

### Competitive repopulation assays

Competitive repopulation assays were carried out as previously described(42). Briefly, BM cells were harvested from both hind limbs of 3-4 donor mice (12-16 weeks old) and mixed with competitive CD45.1 cells at a donor equivalent ratio of 1:1. The mixed BM cells were transplanted into lethally-irradiated recipient CD45.1 mice (10.5 Gy, Cs^137^ source). The level of long-term engraftment was determined by flow cytometry in recipient mice 16 weeks after transplantation using peripheral blood staining with CD45.2, CD45.1, Gr1, B220, Ter119 and CD4 antibodies (Thermo Fisher Scientific, Norcross, GA) or bone marrow c-Kit^+^Lineage^neg^Sca-1^+^ (KLS) cells with CD45.2, CD45.1 along with KLS markers antibodies upon euthanasia.

### Multi-parameter flow cytometry analysis

Bone marrow cells were stained with FITC conjugated lineage antibodies (Gr-1, Mac-1, B220, Ter119, CD3, CD4, CD8a), APC-Cy7-c-Kit, PE-Cy7-Sca-1, eFluor 450-CD48, PerCP-eFluor 710-CD150, eFluor 660-CD34, Alexa Fluor 700-CD16/32, PE-CF594-CD127 (IL7Rα). In some cases, bone marrow cells were stained with Biotin-CD135 (FLK2), then PE-streptavidin along with the antibodies mentioned above. Stained samples were run on FACSymphony A5 (BD Biosciences) and analyzed by FlowJo software (FlowJo, Ashland, OR). The population of cells was defined(27) as CD150^+^CD48^neg^KLS (HSCs), CD150^neg^CD48^neg^KLS (previously referred to as MPP(27)), CD150^neg^CD48^+^KLS (HPC-1) and CD150^+^CD48^+^KLS (HPC-2). When bone marrow cells were stained including CD34 and CD135 antibodies, stem and progenitor cells were defined as from Wilson et al.(28): LT-HSC (CD34^neg^CD48^neg^CD150^+^CD135^neg^KLS), MPP1 (CD34^+^CD48^neg^CD150^+^CD135^neg^KLS), MPP2 (CD34^+^CD48^+^CD150^+^CD135^neg^KLS), MPP3 (CD34^+^CD48^+^CD150^neg^CD135^neg^KLS) and MPP4 (CD34^+^CD48^+^CD150^neg^CD135^+^KLS).

### Cell culture and ELISA

All cell lines were tested for authenticity by short-tandem repeat (STR) analysis in the Emory Integrated Genomics Core Facility and tested mycoplasma negative (PromoKine PCR-based assay kit) prior to cryopreservation of long-term storage vials. EML C1 cells were kindly provided by Schickwann Tsai and were cultured and maintained in IMDM supplemented with 20% (vol/vol) horse serum, 10 ng/mL SCF. Wild-type (WT) and STAT5ab^null/null^ MEF cells were cultured in high-glucose DMEM, 10% FBS, 1x penicillin-streptomycin solution. EML C1 and MEF co-culture experiments were carried out by seeding 0.1 million irradiated (15 Gy, γ-rays ^137^Cs) WT or STAT5ab^null/null^ MEFs per well into 6-well plate 24 hour before, then adding 0.1 million of EML C1 cells per well for co-culturing either 4 or 7 days. Primary murine MSC cultures were carried as described(78). To prevent hematopoietic cell contamination, frequent changes of the medium were done in the first three days followed by lifting the cells with less than a 2 minute treatment with 0.25% trypsin/1 mM ethylenediaminetetraacetic acid (EDTA) and discarding the non-lifted cells. Some STAT5 knockout MSCs were generated from STAT5ab^flox/flox^ mice. At the end of the second week of culture, cells were transduced with a high viral titer Ad-Cre-GFP adenovirus (Baylor College of Medicine, Gene Vector Core). After three weeks of culture, MSCs were checked by flow cytometry for expression of Sca-1(Ly6a) and CD140a. STAT5 deletion was checked by Western blot analysis or real-time PCR. To check for the production of secretory niche factors, wild type or STAT5ab-deficient MEFs or MSCs were cultured. The same number of cells (0.1 million) were seeded in 24-well plate. The second day, fresh DMEM medium with reduced FBS (2%) was changed and the supernatant was harvested 24 hr later for IGF-1 and SDF-1α ELISA assays (R&D Systems, Minneapolis, MN).

### Real-time quantitative reverse transcriptase-polymerase chain reaction (qRT-PCR)

Total cellular RNA was extracted using Trizol reagent (Invitrogen, Carlsbad, CA) according to the manufacturer’s protocol and cDNA was synthesized from 1.0 μg of RNA using the SuperScript III first-strand synthesis system (Thermo Fisher Scientific, Norcross, GA). Real-time PCR was conducted using iQ SYBR Green Supermix (Bio-Rad, Hercules, CA) and amplification was performed on a CFX Touch Real-Time PCR Detection System (Bio-Rad, Hercules, CA) and normalized to glyceraldehyde-3-phosphate dehydrogenase (GAPDH). All PCR primers are from PrimerBank (Harvard University) which has been validated for the real-time PCR.

### Western blotting

Cells were lysed using RIPA lysis and extraction buffer with protease inhibitors and phosphatase inhibitors from Roche (Indianapolis, IN). Protein concentration were determined with the Bio-Rad protein assay (Hercules, CA). Proteins were resolved by 10% SDS-PAGE and transferred to PVDF membrane, and detected by immunoblotting with specific primary antibodies. After blocking in 5% BSA for 1 hour, membranes were incubated in the appropriate antibody overnight and detection was by chemiluminescence (Thermo Fisher Scientific, #34080) or by the Odyssey CLIx imaging system (LI-COR Biosciences). Image Studio v4.0 software was used for densitometry quantification. Antibodies for Western blot were obtained from the following: Primary antibody concentrations were used according to the manufacturers’ suggestions. Secondary antibody combinations used were either IRDye 680RD (925–68070)/IRDye 800CW (925–32210) or IRDye 680RD (925–68071)/IRDye 800CW (925–32211). These Goat anti-Rabbit (LI-COR) antibodies were used at 1:10,000 to 1:20,000 dilution.

### Flow cytometry analyses and cell processing/sorting for single cell RNA sequencing

Bone marrow cells were harvested from the femurs and tibias of 4 Vav1-Cre/+ or Vav1-Cre/+/STAT5ab^flox/flox^ mice (8-12 weeks). The cells were lineage depleted using a mouse lineage cell depletion kit (Miltenyi Biotech, Auburn, CA) and stained with FITC-labeled lineage antibodies (Gr-1, Mac-1, B220, Ter119, CD3, CD4, CD8a), anti-c-Kit-APC, anti-Sca-1-PE-Cy7 (Thermo Fisher Scientific, Norcross, GA), and dead cells were excluded by staining with propidium iodide dye. KLS cells were sorted (sorting purity ∼6% with 605 CD45^neg^ cells among 10008 total sorted KLS cells) using BD FACSAria III Cell Sorter. For sorting bone marrow stromal cells, cells were isolated as described(79). Briefly, mouse bones (tibia and femur) were isolated and cleaned from 6-8 weeks old mice, then were crushed and cut into tiny pieces with sterile scissors, incubated them in 20 ml of preheated DMEM plus 0.2% (wt/vol; 40 mg) collagenase at 110 rpm at 37°C for 1 hour). The cell suspension was filtered through a 70-μm cell strainer and bone fragment was further crushed and washed to liberate cells out of the bone into solution. CD45^+^ cells were depleted with mouse CD45 microbeads from Miltenyi Biotech(Auburn, CA) and stained with anti-Ter119-PE (BD Biosciences, San Jose, CA), anti-CD45-FITC, Anti-CD71-Alexa 700, Anti-Sca-1-PE-Cy7 and Anti-CD140a-APC (Thermo Fisher Scientific, Norcross, GA). CD45^neg^Ter119^neg/low^CD71^neg^Sca-1^+^CD140a^+^ MSCs were sorted using BD FACSAria III Cell Sorter. Sorted cells were immediately processed for single cells RNA sequencing.

### Single-cell RNA sequencing

Tissue derived single cells were loaded onto the 10X Chromium Controller targeting 3000-6000 cells/sample. Single cell capture, barcoding, GEM-RT, clean-up, cDNA amplification and library construction were performed according to the manufacturers’ instructions for v3 chemistry using 10x Genomics Chromium Chip B Single Cell Kit (PN 1000074) and Chromium Single Cell 3’ GEM, Library & Gel Bead Kit v3 (PN 1000092). Final gene expression library pools were sent to Genewiz (South Plainfield, NJ) and sequenced on an Illumina Novaseq6000 instrument with a targeted sequencing depth of 150,000 reads/cell.

### Single-cell transcriptomic data analysis

Raw sequencing reads were processed by Cell Ranger (10x Genomics) to generate gene expression count matrices for every sample (wild type and knockout samples for KLS and MSC respectively). After sequence alignment to GRCm38/mm10 mouse reference genome, UMI (Unique Molecular Identifiers) counts for each gene per cell formed the gene expression count matrices. Based on UMI counts and cell barcodes, numbers of genes and cells for each sample were estimated. Clustering and UMAP visualization based on concatenation of the wild type and knockout samples did not show noticeable batch effect, which was consistent with the fact that the samples were sequenced together. Therefore, subsequent clustering and trajectory analyses were performed based on concatenation of the samples, without applying any scRNA-seq data integration workflow. Data for KLS and MSC were analyzed separately. Implemented using the Seurat package in R, the clustering workflow started with library size normalization and log transformation, followed by highly variable feature selection, principle component analysis and community detection, with clusters visualized using UMAP (Uniform Manifold Approximation and Projection). After cell clustering, differential gene expression analysis was performed using Wilcoxon rank sum test between each cell cluster against all other cells to obtain marker genes for interpreting the cell clusters. Cell clusters were annotated by manually examining the marker genes derived from the data, based on prior knowledge in the literature. Implemented using the Scanpy package in Python, the trajectory analysis was performed using the PAGA (partition-based graph abstraction) algorithm. The cell clusters generated in the Seurat clustering analysis served as the input of cell clusters to the PAGA algorithm, so that nodes in the graph abstraction of PAGA can be directly mapped to the cell clusters in the clustering analysis.

### Gene-set enrichment analysis (GSEA)

Several GSEA algorithms were used to identify aberrant pathway and biological changes in the data sets by examining differentially-expressed genes comparing wild-type and knockout bone marrow cells. From those lists of the top upregulated and top downregulated genes, the following algorithms are presented in the figures: GO Biological Processes 2021, MSigDB Hallmark 2020, and Panther 2015. Panther was used for all other analyses where the data are described in the text only and not part of a specific Table or Figure.

### Co-expression algorithms

Maximum Mean Discrepancy (MMD) measures the distance between two distributions. In our analysis, scRNA-seq datasets undergo conventional pre-processing steps including gene and cell filtering, high-mitochondrial cells removal, library-size normalization, and highly-variable gene selection, principal component analysis, followed by tSNE for dimension reduction. In the tSNE space, given a gene of interest, coordinates of cells expressing the gene are extracted. For another gene, coordinates of cells expressing the other gene are also extracted, separately. To quantify the co-expression of those two genes, the two sets of coordinates are used as input for the MMD algorithm to quantify the distance between the two distributions. Smaller MMD distances indicate two distributions are similar, hence stronger co-expression between the two genes. On the contrary, larger MMD distances indicate two distributions are dissimilar, hence weaker co-expression between the two genes. The logarithm of the MMD scores were used for subsequent analysis.

### Statistical analyses

All data were derived as a result of three or more independent experiments, unless stated otherwise. Student’s two tailed t-test was used to calculate p-values and values less than 0.05 were considered to be significant. Throughout all figures the following nomenclature was used to indicate the level of statistical significance: * P≤0.05; ** P≤0.01; *** P≤0.001.

### Materials and data availability

The newly generated mouse models described here will not all be sustained long-term in the lab. However, all mouse strains used are commercially available, except the Cdh5-Cre mice. All strain numbers from JAX have been provided. Antibodies used are listed in **Table S5** and DNA oligonucleotide sequences used for qRT-PCR are listed in **Table S6**. EML c1 cells are commercially available. STAT5ab knockout MEFs were generated in the Bunting lab and will be provided as needed. Newly generated scRNAseq data has been submitted to GEO with accession # GSE214857. The MMD co-expression code used in this manuscript was submitted to the GitHub repository. Publicly available scRNAseq data was obtained from the following sources: NCBI Gene Expression Omnibus, GSE128423, GSE108892, GSE132151, GSE100428, and GSE81682.

## Acknowledgements

This work was funded by the I3 Rapid RFP Emory SOM/Georgia Tech Computational and Data Analysis to Advance Single Cell Biology Research Award (K.B. and P.Q.); Aflac Cancer and Blood Disorders Center Pilot Grant (K.B.), NIHR01DK059380 (K.B.), Aflac Cancer & Blood Disorders Center of Children’s Healthcare of Atlanta and Emory University School of Medicine (K.B.) We acknowledge the generous support of the Emory + Children’s Pediatric Research Center and the Emory/Pediatrics Winship Flow Cytometry Core. Additional technical support was provided by Emory Integrated Genomics core for single cell RNAseq analysis.

## Competing Interests

Dr. Bunting owns a consulting company in the State of Georgia called Valhalla Scientific Editing Service, LLC.

---

**Fig. S1.**
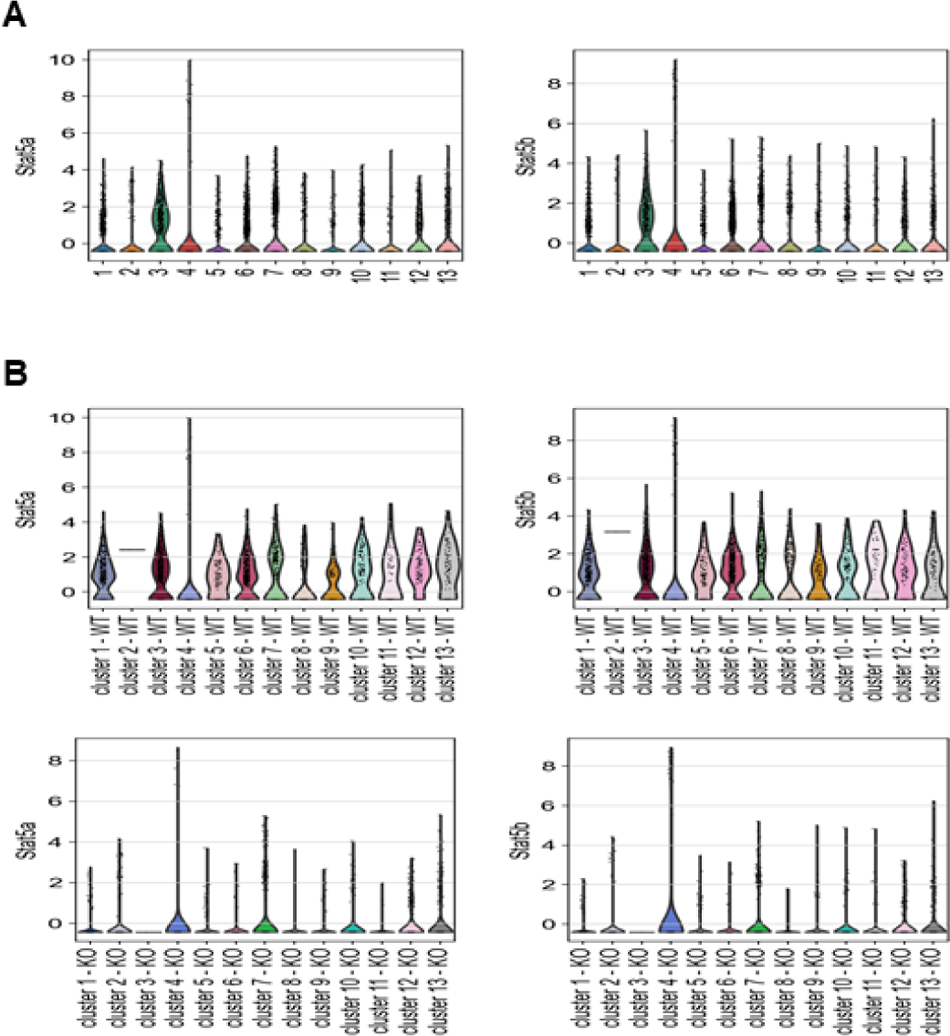
Expression levels of Stat5ab in all 13 clusters from sorted KLS cells revealed as determined by single cell RNA seq analysis. (A). Violin plots showing the distributions of Stat5a and Stat5b expression in each cluster. (B). Violin plots showing differential expression of Stat5a and Stat5b in each cluster between WT and KO.

**Fig. S2.**
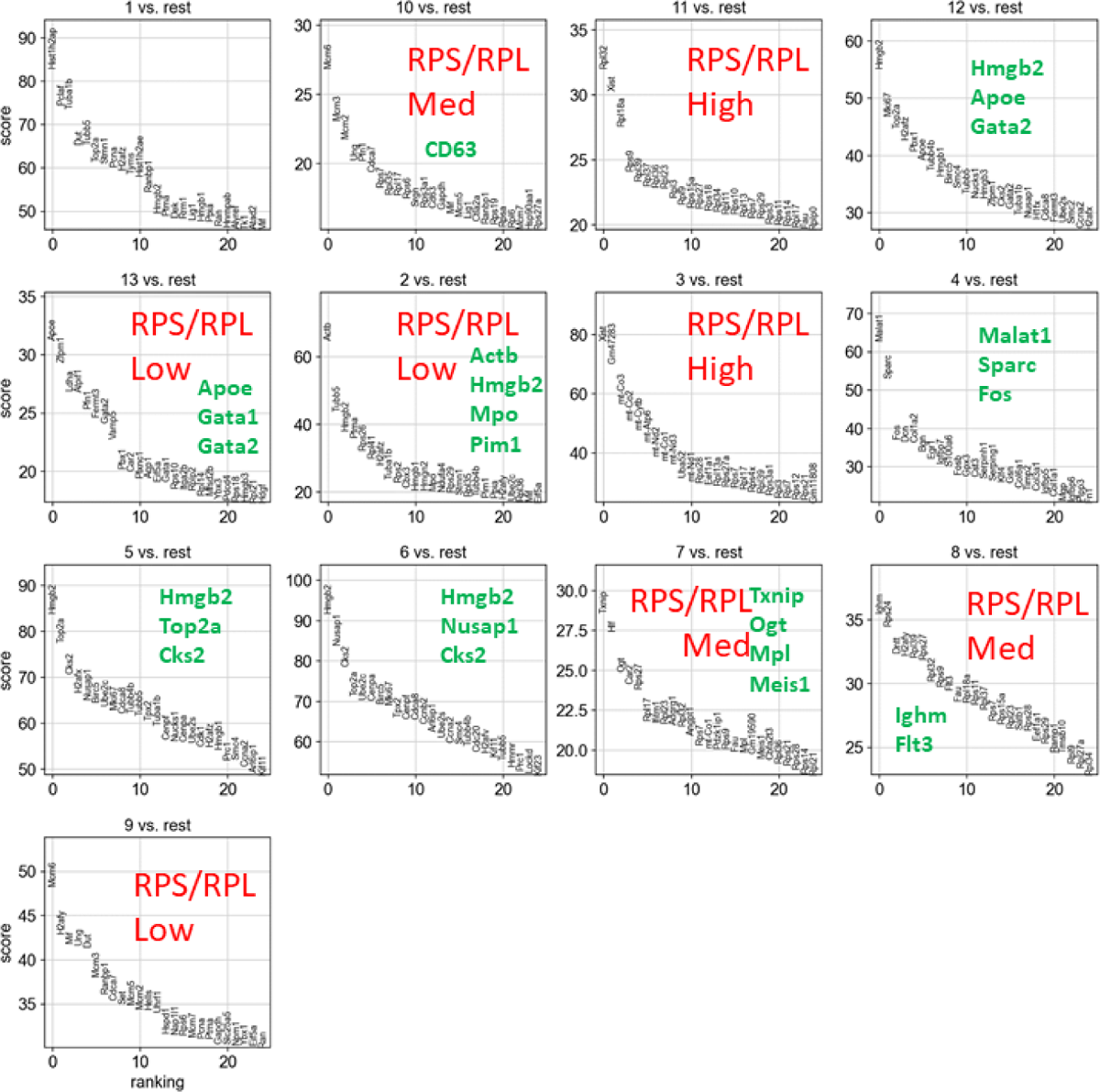
Top differentially expressed genes defining the 13 clusters. The t-test scores of each cluster vs the rest are shown for the top 25 differentially expressed genes for each cluster. High levels of myeloid priming were observed with the trajectory going through myeloid-biased MPP3-like cells. Peripheral blood hematology analysis of Vav1-Cre/+STAT5ab^fl/fl^ mice showed decreased granulocyte-monocyte-lineage cells (increased percentage), decreased hematocrit and red blood cell count, and as expected dramatically decreased overall white blood cell count and lymphocyte count (**Table 1**).

**Fig. S3.**
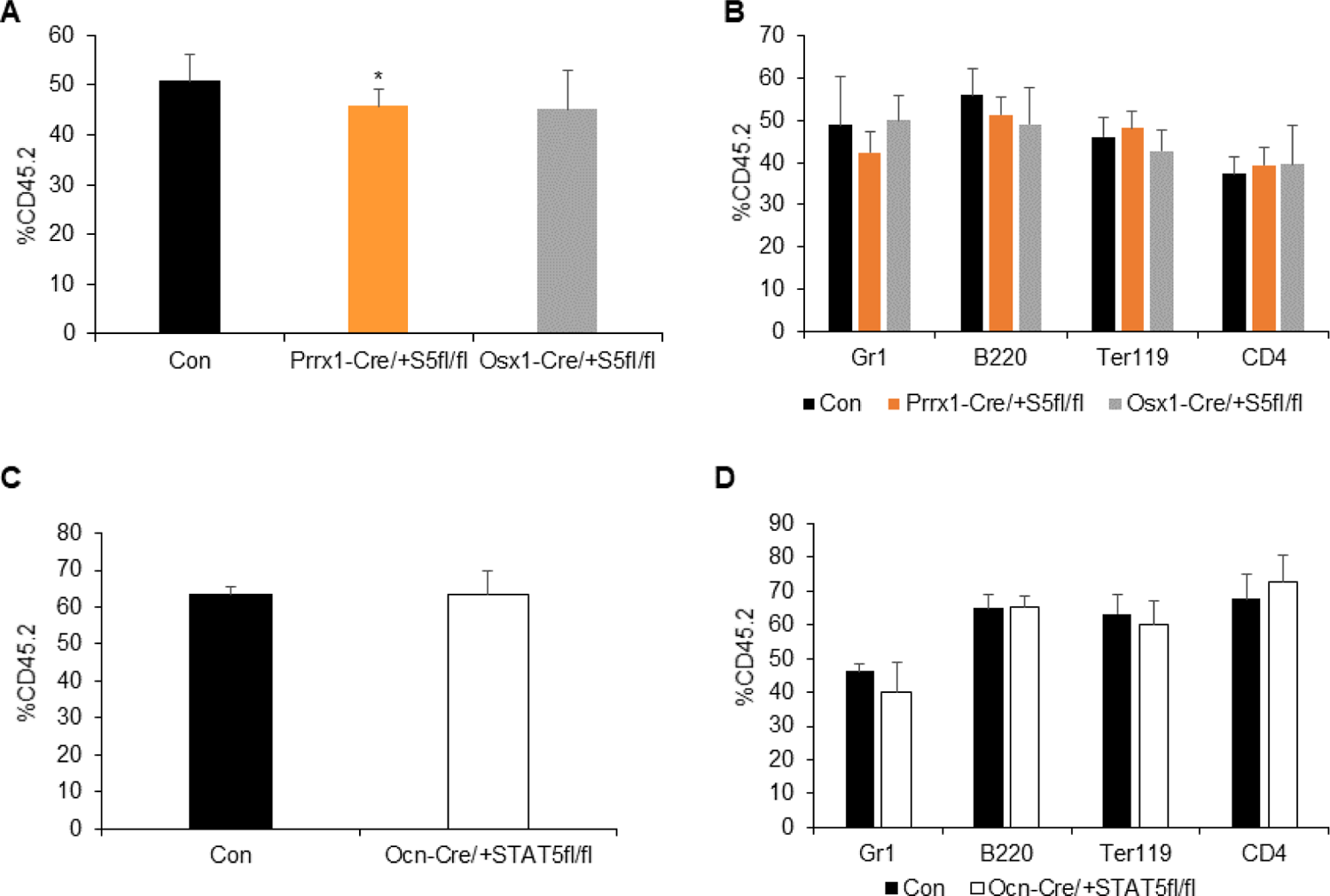
Prrx1-Cre/+, Osx1-Cre/+, and Ocn-Cre/+ deletion of STAT5ab does not alter the bone marrow HSC competitive repopulation activity. Bone marrow cells from 3-4 donor mice were collected from each group, mixed 1:1 with wild type competitor (CD45.1) and transplanted into lethally-irradiated Boy J mice (CD45.1) for competitive repopulation analysis. Recipient mice were bled 16 weeks later for flow cytometry analysis. The genotype STAT5ab^fl/fl^ is also represented here as S5fl/fl or STAT5^fl/fl^ because of space. (A, B) Competitive bone marrow repopulation assay for Prrx-1/+STAT5ab^fl/fl^ and Osx1-Cre/+STAT5ab^fl/fl^ mice. The overall donor percentage of CD45.2 positive in the peripheral blood is shown in (A) and the result of the multi-lineage analysis is presented in (B). Results are from the average two independent experiments with 5 mice per group in each experiment. (C, D). Competitive bone marrow repopulation assay for Ocn-Cre/+STAT5ab^fl/fl^ mice. Overall donor engraftment is in (C) and multi-lineage donor engraftment is in (D). Results are from one experiment with 5 mice per group.

**Fig. S4.**
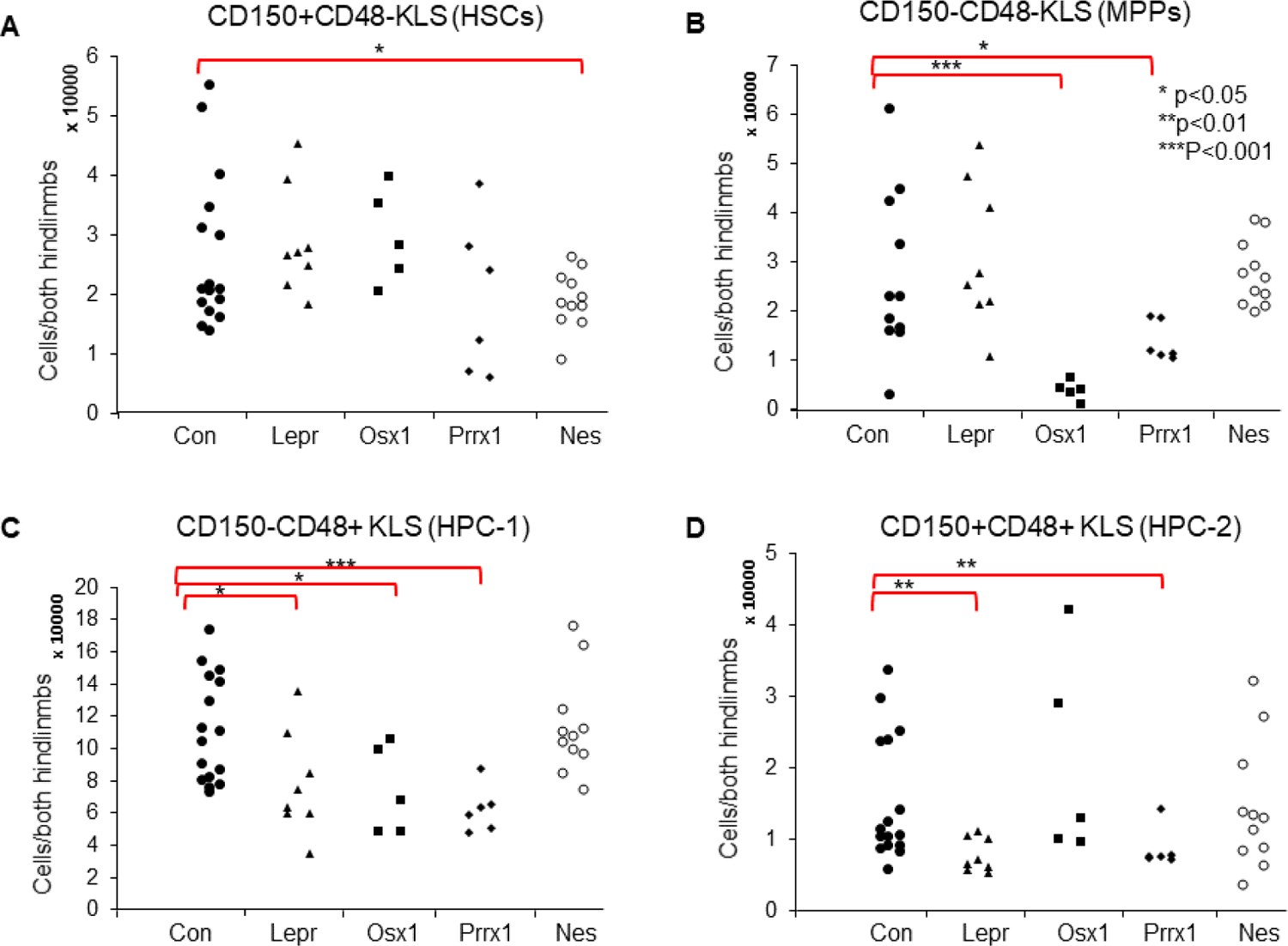
Conditional deletion of STAT5ab with Nes-Cre, Lepr-Cre, Prrx1-Cre, and Osx1-Cre altered the number of stem and progenitors in the bone marrow. Bone marrow cells from Nes-Cre/+STAT5ab^fl/fl^, Lepr-Cre/+STAT5ab^fl/fl^, Prrx1-Cre/+STAT5ab^fl/fl^, Osx1-Cre/+STAT5ab^fl/fl^ and control Vav1-Cre/+ were assayed by multi-parameter flow cytometry to quantitate the number of stem and progenitors. BM cells were stained with antibodies against lineage markers, c-Kit, Sca-1, IL7R, and SLAM marker antibodies as well as CD150, CD48. KLS cells were further subdivided into four stem and progenitor population based on the staining of CD150 and CD48. The absolute number of stem and progenitor cells are presented in dot plots. (A). CD150^+^CD48^neg^KLS (enriched for LT-HSC). (B). CD150^neg^CD48^neg^KLS (MPPS). (C). CD150^neg^CD48^+^KLS (HPC-1). (D). CD150^+^CD48^+^KLS (HPC-2). STAT5ab deletion with Lepr-Cre reduced restricted hematopoietic progenitor cells (HPC-2 and HPC-1(27)) with Osx1 and Prrx1-Cre on the reduction of restricted progenitors and MPPs, while only Nes-Cre knockout STAT5 had significantly reduced HSCs. Those results are consistent with a more progenitor supporting role for STAT5ab throughout the mesenchymal osteo/chondro-genic lineages and Nes expression being promiscuous in HSCs. Ocn-Cre driven deletion of STAT5ab did not have a noticeable effect (Fig. S5A/B). Tamoxifen-induced Nes-CreERT2/+ to delete STAT5ab led to reduced MPPs but not HSCs (Fig. S5C/D).

**Fig. S5.**
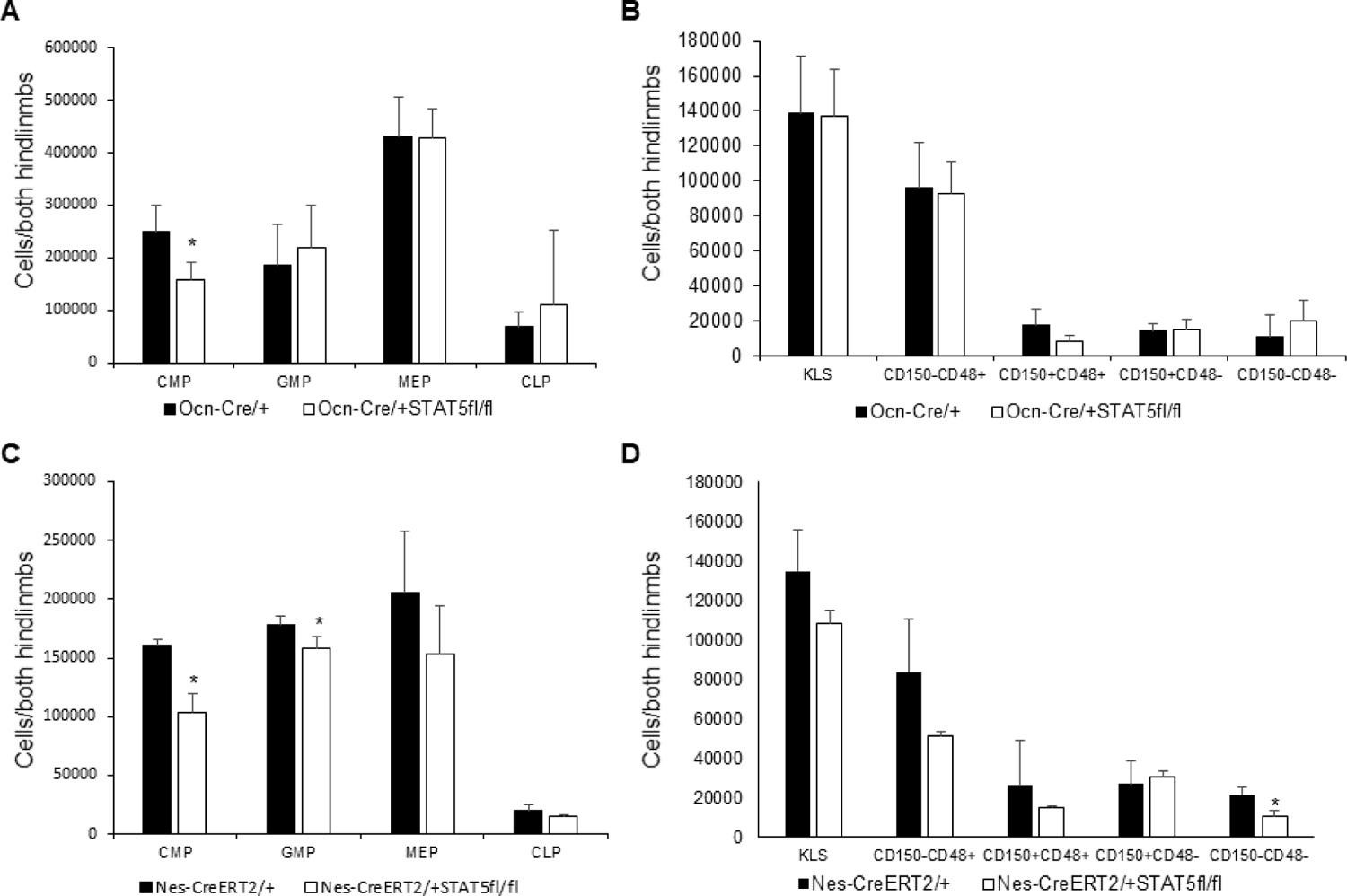
Conditional deletion of STAT5ab with Ocn-Cre and Nes-CreERT2 has minimal effects on bone marrow hematopoietic stem and progenitors. BM cells were stained with antibodies against lineage markers, c-Kit, Sca-1, CD150, CD48, CD16/32, CD34, and IL7R followed by flow cytometry. The genotype STAT5ab^fl/fl^ is also represented here as STAT5^fl/fl^ because of space. (A, B) The absolute number of bone marrow stem and progenitor cells in Ocn-Cre/+STAT5ab^fl/fl^ and its control Ocn-Cre/+ mice (A) Lineage-committed progenitors including CMP, GMP, MEP and CLP. (B) Stem and progenitor cells based on the KLS subdivided by CD150 and CD48 staining (N=5 Ocn-Cre/+, n=4 for Ocn-Cre/+STAT5ab^fl/fl^ mice). (C, D) The absolute number of bone marrow stem and progenitor cells in Nes-CreERT2/+STAT5ab^fl/fl^ and its control Nes-CreERT2/+ mice. Nes-CreERT2/+STAT5ab^fl/fl^ and its control Nes-CreERT2/+ mice (12-16 weeks old) were treated with 75 mg tamoxifen/kg body weight via i.p. injection for a total of 5 consecutive days. Mice were analyzed two weeks after the last treatment. (C) Lineage committed progenitors including CMP, GMP, MEP and CLP. (D) Stem and progenitor cells gated on KLS markers then subdivided by CD150 and CD48 staining (n=4 for Nes-CreERT2/+, n=3 for Nes-CreERT2/+STAT5ab^fl/fl^ mice). Additionally, Tamoxifen-induced Cdh5-Cre did not have effects on either progenitors or HSCs (**Fig. S6**) indicating that the role for STAT5ab in stroma is in a perivascular cell type but not in the vasculature itself.

**Fig. S6.**
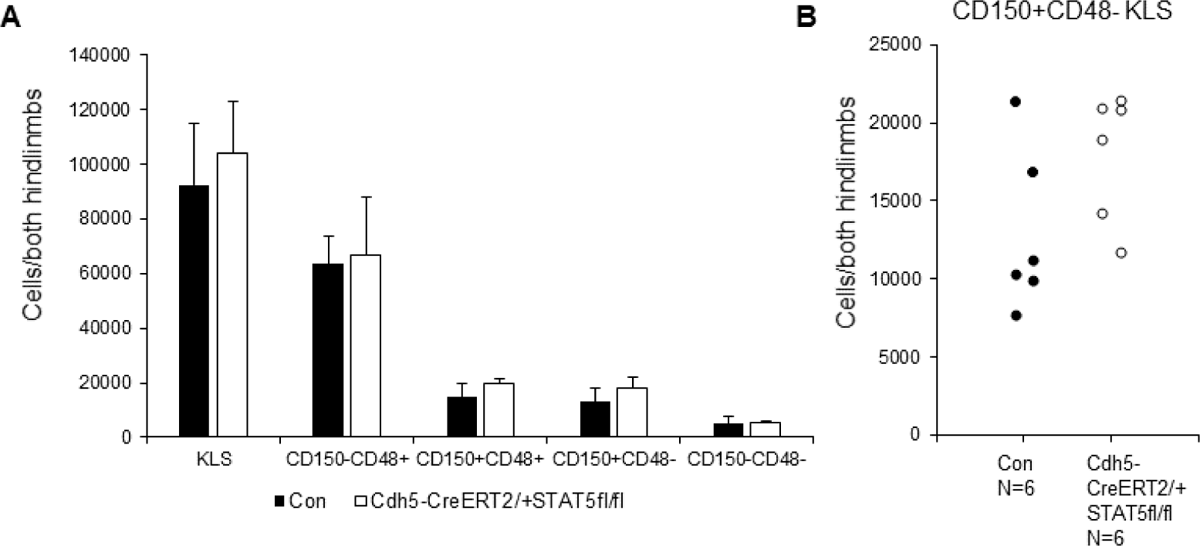
Conditional deletion of STAT5 with Cdh5-CreERT2 has no effect on bone marrow hematopoietic stem and progenitors. Tamoxifen treatment was followed by flow cytometry analysis using the same approach as in Fig S5. The genotype STAT5ab^fl/fl^ is also represented here as STAT5^fl/fl^ because of space. (**A**) Lineage-committed progenitors including CMP, GMP, MEP and CLP were assessed by flow cytometry. (**B**) Stem and progenitor cells based on KLS markers were further subdivided by CD150 and CD48 staining (n=6 from each group).

**Fig. S7.**
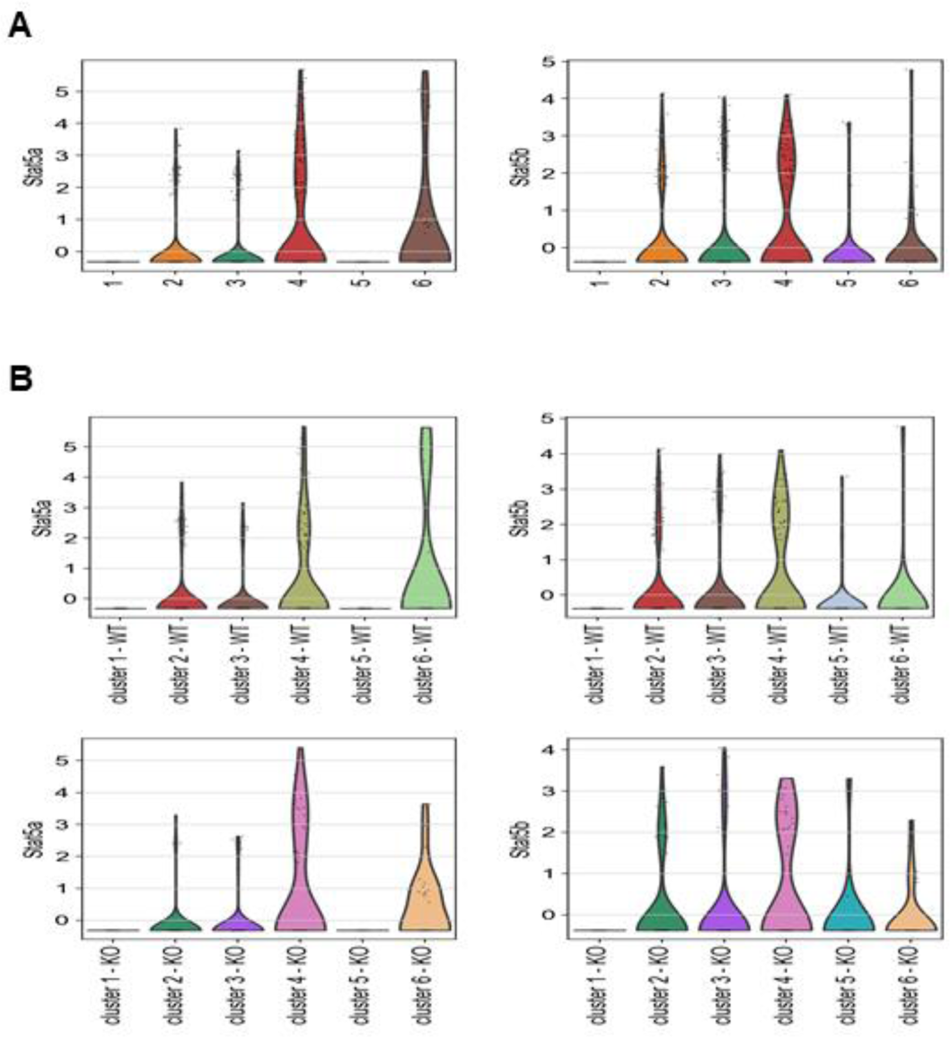
Expression of Stat5ab in 6 clusters defined from the MSC stromal cell population. (**A**). Violin plots showing the distributions of Stat5a and Stat5b expression in each cluster. (**B**). Violin plots showing differential expression of Stat5a and Stat5b in each cluster between WT and KO.

**Fig. S8.**
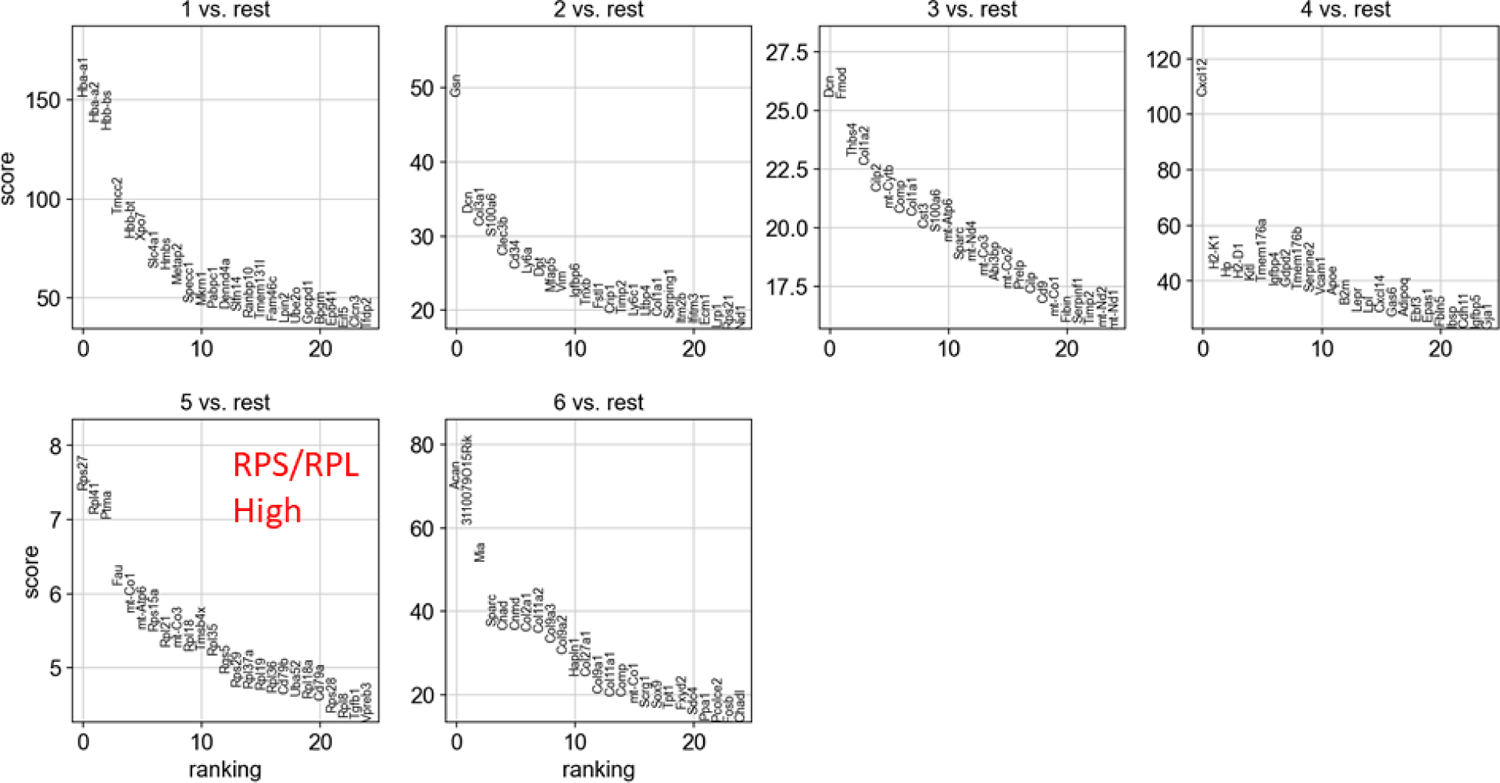
Top differentially expressed genes defining the 6 clusters. The t-test scores of each cluster vs the rest are shown for the top 25 differentially expressed genes for each cluster.

**Fig. S9.**
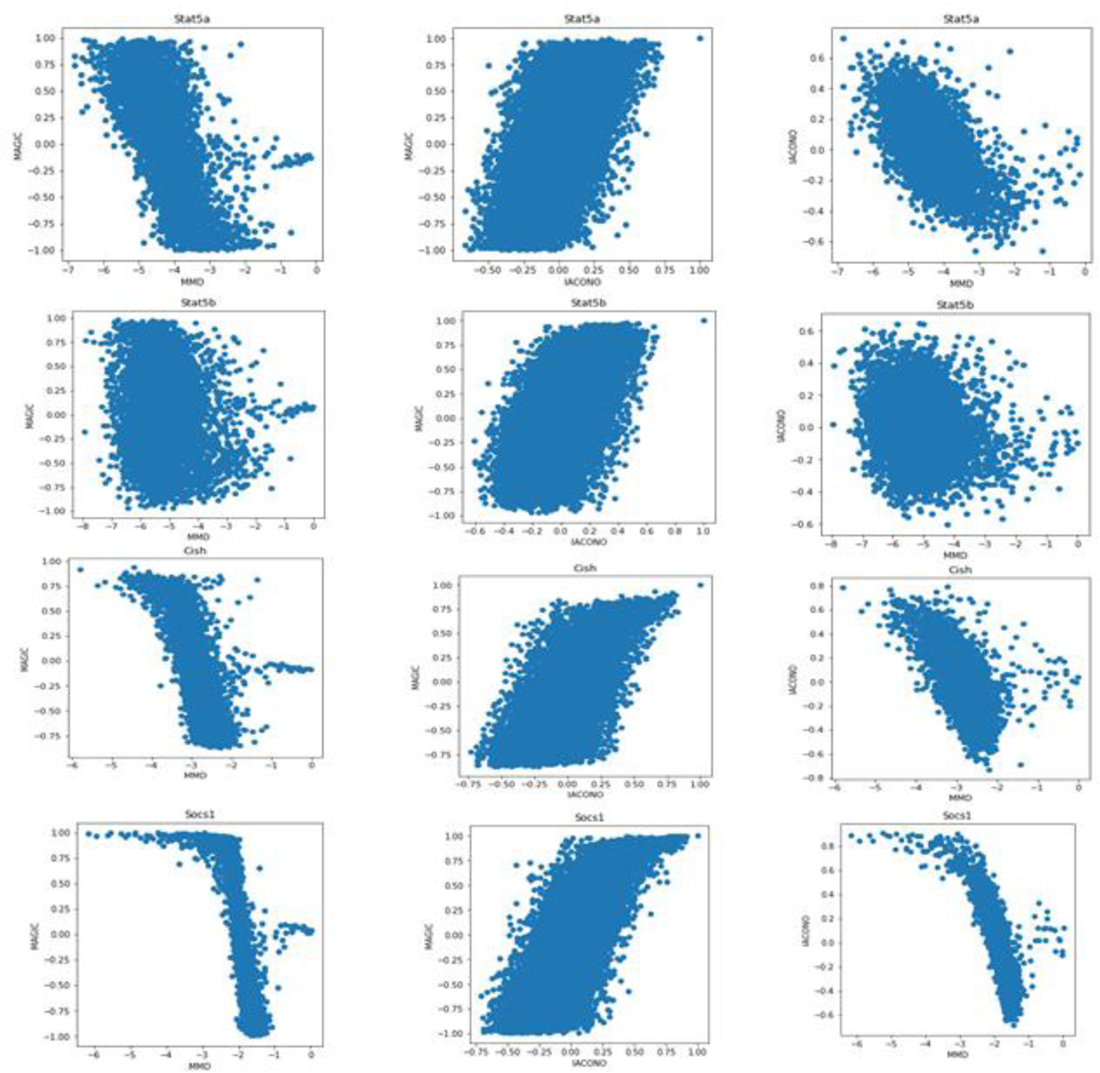
Comparisons of gene co-expression scores of three different methods (MMD, Iacono, MAGIC) on Lepr^+^ cells from the dataset of Scadden. Each row represents a different target gene of interest (Stat5a, Stat5b, Socs1, Cish). Co-expression scores between one target gene and all other genes are computed using three different methods. The three panels in the same row show pairwise comparisons of the co-expression scores from the three methods. In general, MAGIC and Iacono are positively correlated with each other, and MAGIC and Iacono are negatively correlated with MMD scores, which are all expected. The general correlations indicate that the three methods captured similar trends of gene-gene co-expression. The first panel in the last row shows co-expression scores between Socs1 and other genes computed by MAGIC and MMD. In this panel, genes that received low MMD scores all received high MAGIC scores, where both methods suggested strong correlations with Socs1. In contrast, for genes that received high MAGIC scores, their MMD scores were quite variable. In other words, co-expressed genes implicated by MMD were agreeable by MAGIC, but the reverse was not true. Similar pattern was observed in several other panels. Therefore, MMD seems to be more robust and conservative than MAGIC in identifying co-expressed genes.

**Fig. S10.**
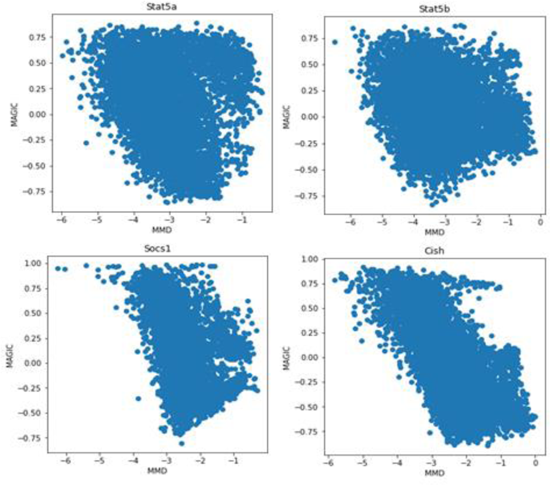
Comparisons of gene co-expression scores of two different methods (MMD and MAGIC) on Nes^+^ cells from the dataset of Scadden. Each panel represents a different target gene of interest (Stat5a, Stat5b, Socs1, Cish). Co-expression scores between one target gene and all other genes are computed using two different methods, and the scores are visualized in one scatter plot. Scores from MMD and MAGIC were in general negatively correlated, which was expected and suggested that the two methods captured a similar co-expression trend in the data. The first panel in the second row shows co-expression scores between Socs1 and other genes computed by MAGIC and MMD. Genes that received low MMD scores all received high MAGIC scores. In contrast, for genes that received high MAGIC scores, their MMD scores were quite variable. Therefore, based on Nes^+^ cells, MMD also seems to be more robust and conservative than MAGIC in identifying co-expressed genes.

**Fig S11.**
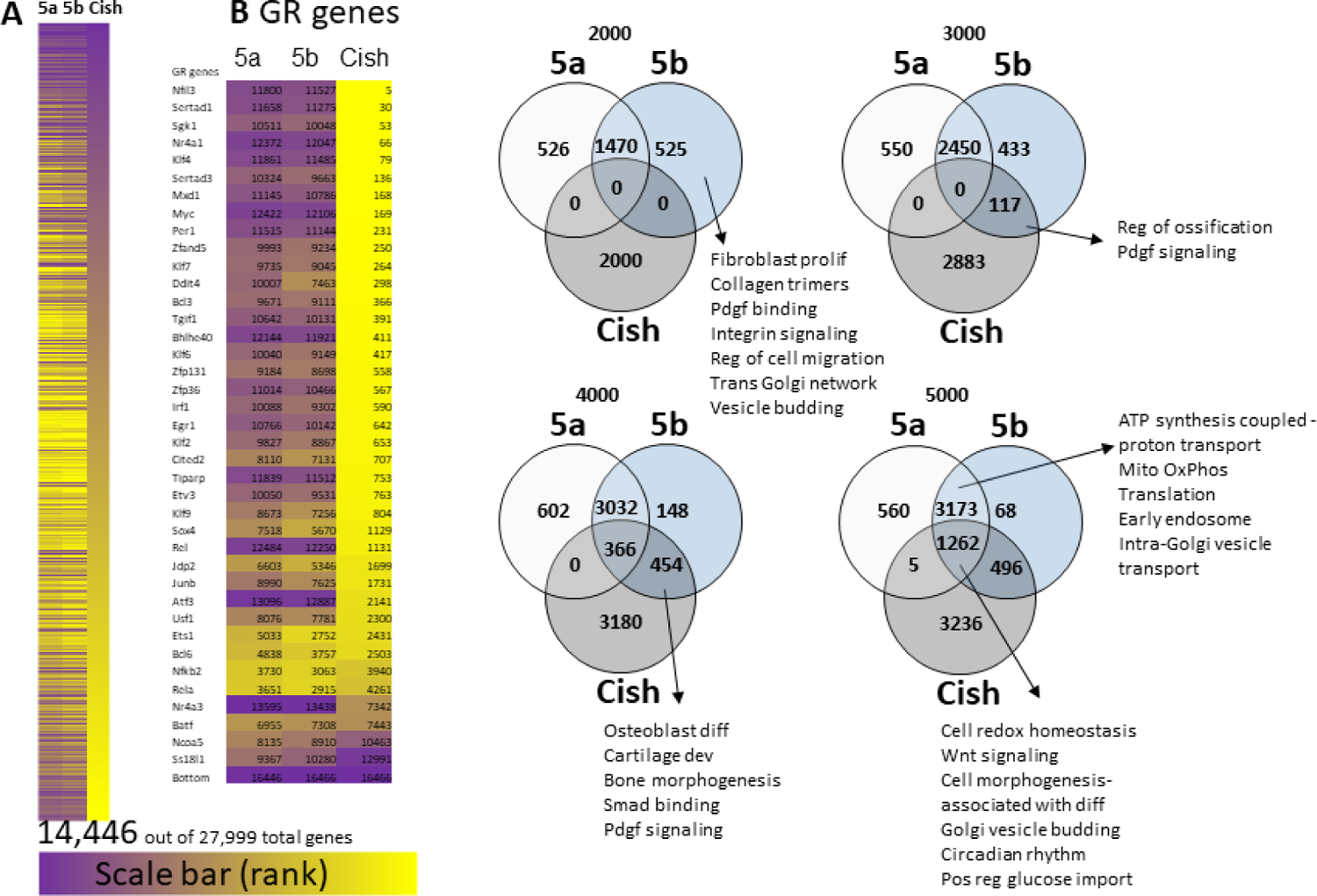
Stat5a, Stat5b, and Cish gene co-expression using Lepr-MSC (cluster 1) from the dataset of Scadden. (**A**). Stat5a, Stat5b, Cish co-expression was analyzed by MMD method using Lepr-MSC (cluster 1) from the dataset of Scadden and ranked by Cish co-expression scores. (**B**). Glucocorticoid response genes have high Cish co-expression scores and are inversely related to Stat5a or Stat5b co-expression scores. (**C**). Venn diagram for the top ranked (top 2000, 3000, 4000 and 5000) Stat5a, Stat5b and Cish co-expression genes and the enriched group functions as defined by GSEA.

**Fig S12.**
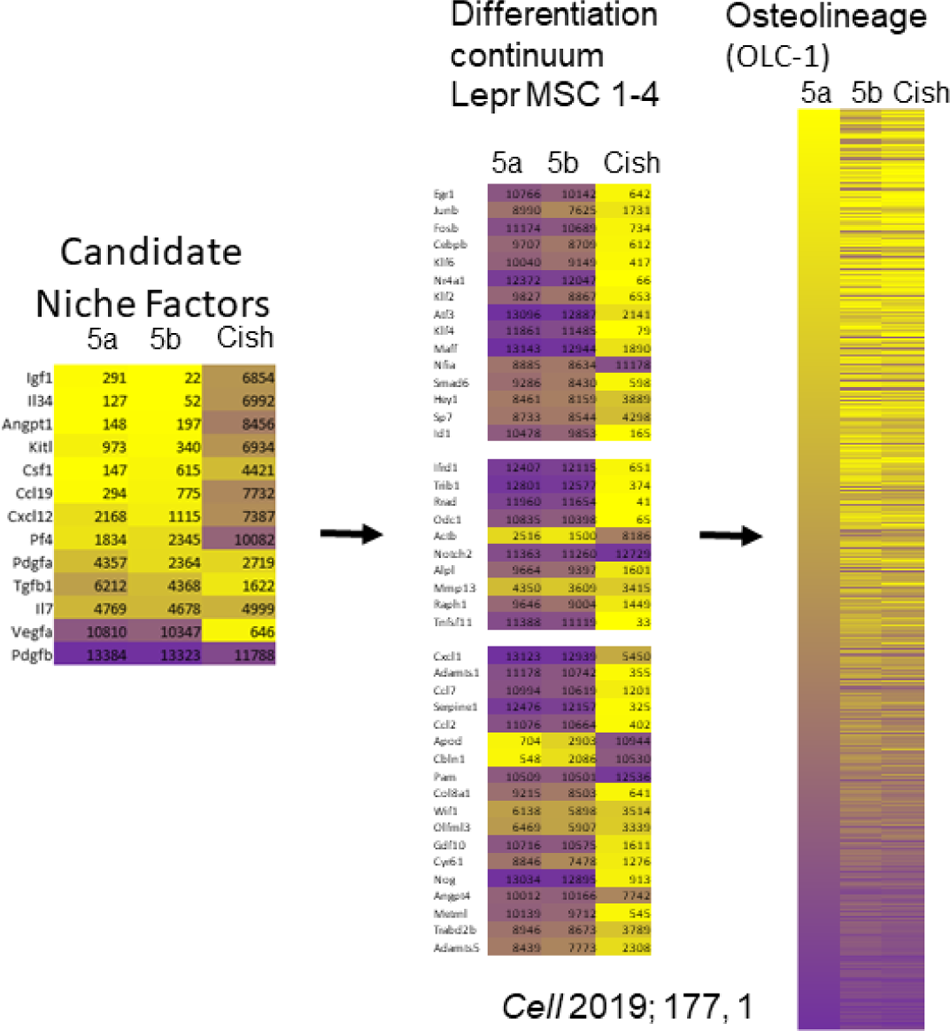
MMD co-expression identifies genes for candidate HSC niche factors and MSC differentiation/glucocorticoid response genes from the dataset of Scadden. Stat5a and Stat5b co-expression in the Lepr-MSC cluster of Scadden was determined by MMD. The highest ranked genes with the most co-expression have the smaller numbers and the lighter (more yellow) color. The lower ranked genes with the least co-expression have the higher numbers and the darker (more purple) color.

**Fig S13.**
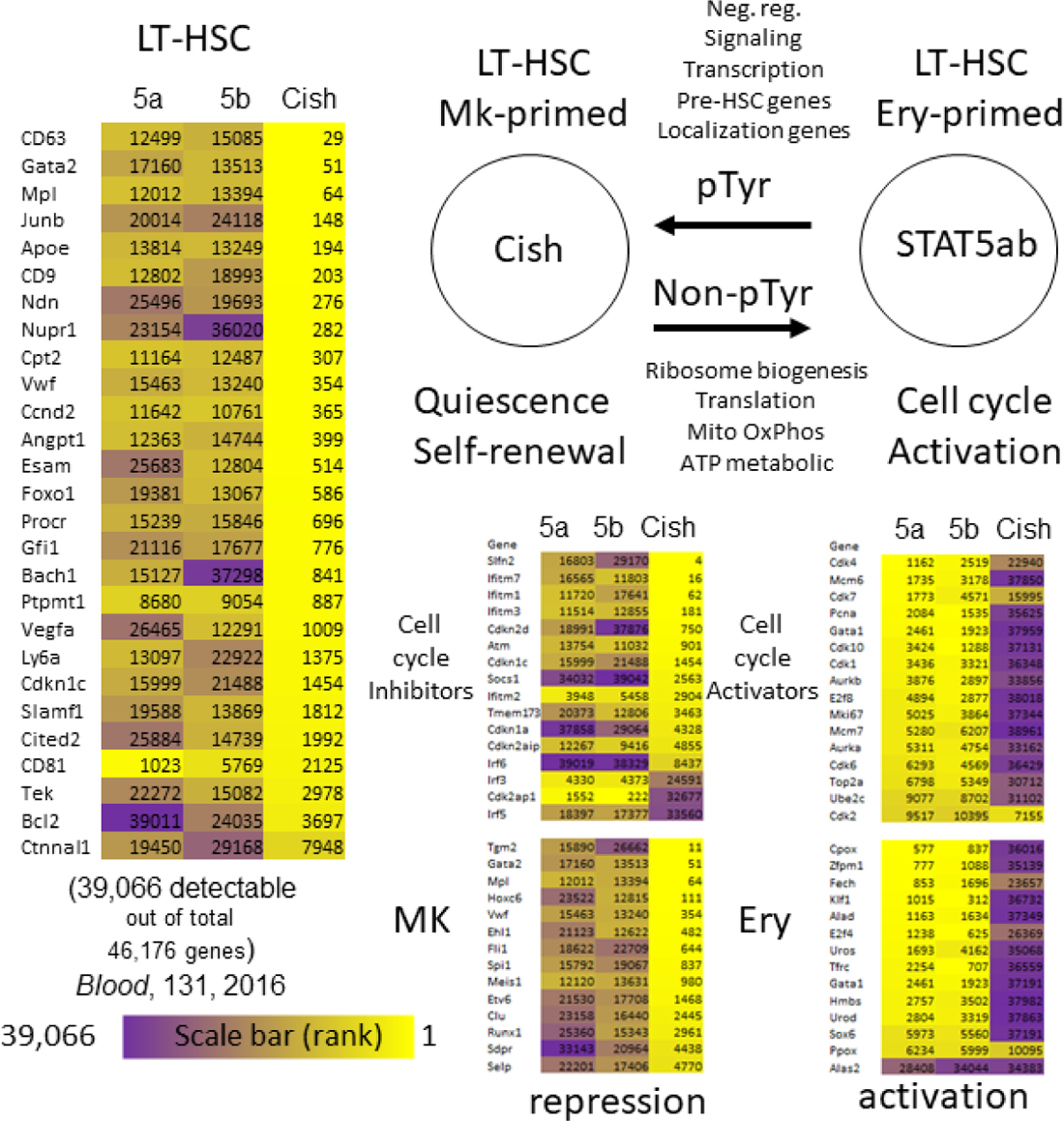
MMD co-expression identifies gene signatures associated with HSC self-renewal vs. lineage commitment decisions in the dataset of Gottgens. Stat5a, Stat5b, and Cish gene co-expression scores were generated from MMD using the LT-HSC cluster from the dataset of Gottgens. The highest ranked genes with the most co-expression have the smaller numbers and the lighter (more yellow) color. The lower ranked genes with the least co-expression have the higher numbers and the darker (more purple) color. A reciprocal gene expression signature was found between STAT5ab and Cish, indicating that the dynamic equilibrium between the transcription factor and its faithful negative feedback regulator target gene is tightly associated with molecular pathways responsible for cell cycle status and Mk/Ery differentiation decisions. MMD comparison of 3 different publicly available data sets (MSC) or 2 different publicly available data sets and our own (HSC) showed overlap among gene sets within the top 5000 most co-expressed genes from independent studies of MSCs or HSCs (**Fig. S14**).

**Fig. S14.**
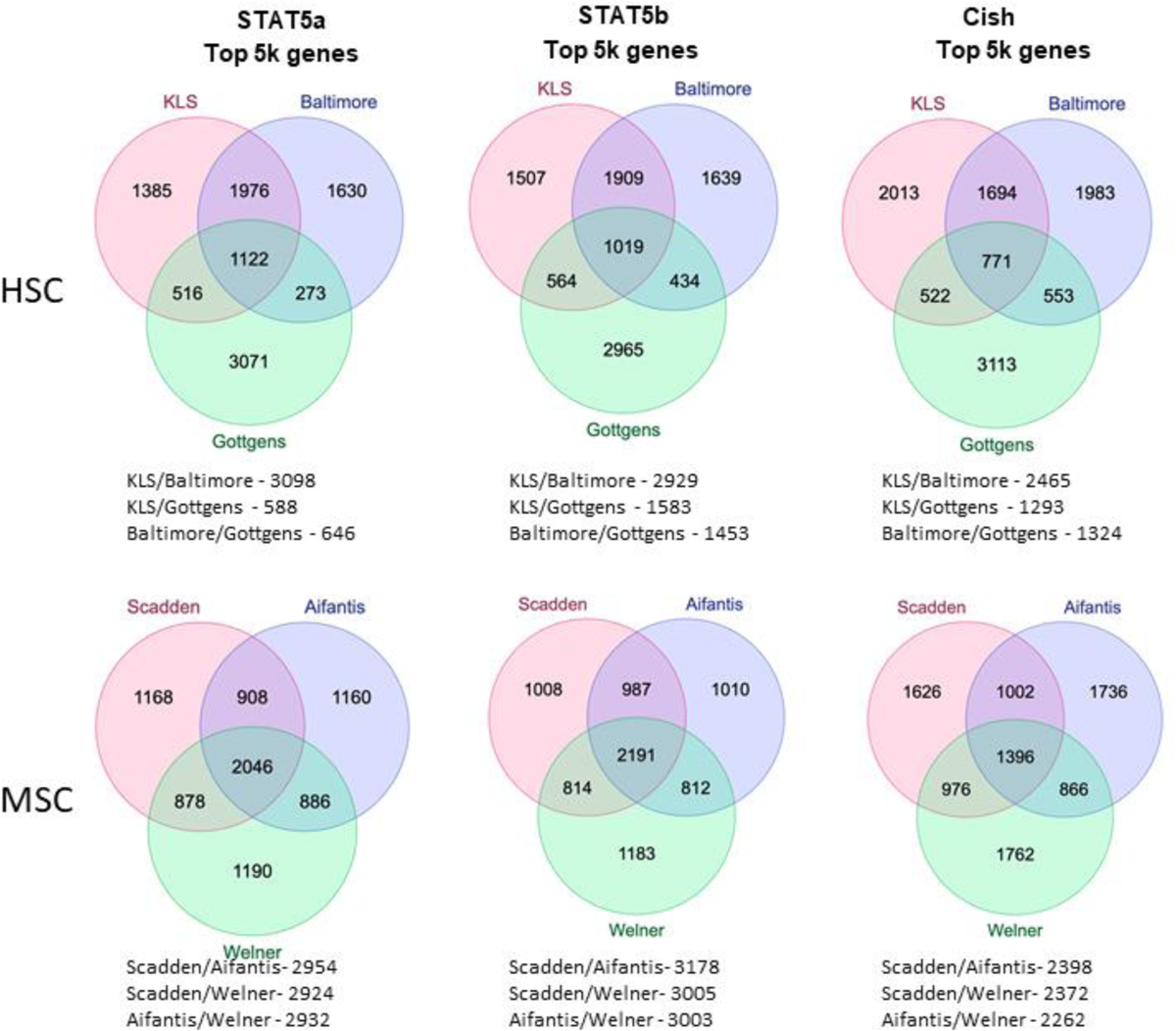
Comparison of gene lists generated by MMD scoring analysis from three different HSC and three different MSC data sets. Venn diagrams capture the overlap between the three different datasets that include LT-HSC clusters. Shown are the top 5000 genes within each cluster and the number of genes that overlap for Stat5a, Stat5b, Cish, and Socs1. Similarly, MSCs were analyzed to compare the overlap between three different datasets that include Lepr-MSC clusters. Shown are the top 5000 genes within each cluster and the number of genes that overlap for Stat5a, Stat5b, Cish, and Socs1. The Scadden dataset was used to interrogate Lepr-MSCs (**Fig. 8A**) and our dataset was used to interrogate HSCs (**Fig. 8B**) respectively. Comparisons were made with both versus our wild-type and STAT5ab knockout MSC and HSC clusters for maximum validation. Combined STAT5ab knockout and wild-type co-expression analyses identified common STAT5ab/Cish regulation of MSC/HSC differentiation priming (**Fig. 8C**) with no distinct differences between STAT5a or STAT5b. Overall, these computational analyses suggest that the STAT5ab-mediated gas/brakes driving MSC/HSC differentiation priming are conserved in two heterotypic bone marrow cell types which cross-talk to regulate normal hematopoiesis.

**Table S1.** GO Biological Process 2021

| Name | P-value | Adjusted p-value |
| --- | --- | --- |
| SRP-dependent cotranslational protein targeting to membrane (GO:0006614) | 1.722e-46 | 4.707e-43 |
| cotranslational protein targeting to membrane (GO:0006613) | 2.083e-45 | 2.847e-42 |
| cytoplasmic translation (GO:0002181) | 1.525e-42 | 1.042e-39 |
| nuclear-transcribed mRNA catabolic process, nonsense-mediated decay (GO:0000184) | 6.138e-44 | 5.593e-41 |
| protein targeting to ER (GO:0045047) | 1.061e-41 | 5.801e-39 |
| peptide biosynthetic process (GO:0043043) | 6.941e-37 | 2.711e-34 |
| nuclear-transcribed mRNA catabolic process (GO:0000956) | 6.692e-37 | 2.711e-34 |
| translation (GO:0006412) | 4.702e-33 | 1.607e-30 |
| rRNA processing (GO:0006364) | 4.547e-24 | 1.036e-21 |
| cellular macromolecule biosynthetic process (GO:0034645) | 1.195e-29 | 3.631e-27 |

**Gene set enrichment analysis (GSEA) using Gene Ontology (GO) biological process 2021 with top differentially expressed genes in KLS cluster 7.**

**Table S2.**
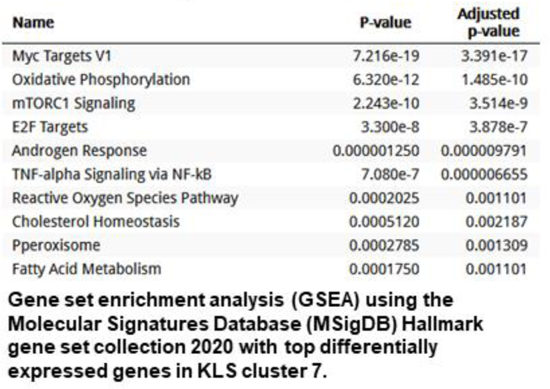
MSigDB Hallmark 2020

| **Name** | **P-value** | **Adjusted p-value** |
| --- | --- | --- |
| Myc Targets V1 | 7.216e-19 | 3.391e-17 |
| Oxidative Phosphorylation | 6.320e-12 | 1.485e-10 |
| mTORC1 Signaling | 2.243e-10 | 3.514e-9 |
| E2F Targets | 3.300e-8 | 3.878e-7 |
| Androgen Response | 0.000001250 | 0.000009791 |
| TNF-alpha Signaling via NF-kB | 7.080e-7 | 0.000006655 |
| Reactive Oxygen Species Pathway | 0.0002025 | 0.001101 |
| Cholesterol Homeostasis | 0.0005120 | 0.002187 |
| Pperoxisome | 0.0002785 | 0.001309 |
| Fatty Acid Metabolism | 0.0001750 | 0.001101 |

**Gene set enrichment analysis (GSEA) using the Molecular Signatures Database (MSigDB) Hallmark gene set collection 2020 with top differentially expressed genes in KLS cluster 7.**

**Table S3.** Panther 2015

| Name | P-value | Adjusted p-value |
| --- | --- | --- |
| Glycolysis | 0.0001235 | 0.002223 |
| Cytoskeletal regulation by Rho GTPase | 0.000007902 | 0.0004267 |
| Cholesterol biosynthesis | 0.001974 | 0.01917 |
| Pentose phosphate pathway | 0.01580 | 0.09483 |
| Ubiquitin proteasome pathway | 0.0006587 | 0.008892 |
| Integrin signalling pathway | 0.00004450 | 0.001201 |
| Huntington disease | 0.002130 | 0.01917 |
| Parkinson disease | 0.003861 | 0.02979 |
| Adenine and hypoxanthine salvage pathway | 0.1189 | 0.2919 |
| JAK/STAT signaling pathway | 0.04654 | 0.1753 |

**Gene set enrichment analysis (GSEA) using the PANTHER (Protein Analysis Through Evolutionary Relationships) Classification System 2015.**

**Table S4.** Peripheral blood hematology of Vav1-Cre/+, Prrx1-Cre/+STAT5ab^fl/fl^, Osx1-Cre/+, Ocn-Cre/+STAT5ab^fl/fl^ mice

|  | Vav1-Cre/+<br>(n=7) | Prrx1-Cre/+<br>STAT5fl/fl<br>(n=8) | Osx1-Cre/+<br>STAT5 fl/fl<br>(n=9) | Ocn-Cre/+<br>STAT5fl/fl<br>(n=6) |
| --- | --- | --- | --- | --- |
| WBC (10<sup>3</sup>/μl) | 10.0 ± 1.5 | 7.7 ± 1.7* | 9.7 ± 2.6 | 11.3 ± 1.4 |
| LYM (10<sup>3</sup>/μl) | 8.5 ± 1.1 | 6.2 ± 1.4** | 8.4 ± 2.3 | 9.4 ± 1.2 |
| MONO (10<sup>3</sup>/μl) | 0.5 ± 0.1 | 0.5 ± 0.1 | 0.5 ± 0.1 | 0.6 ± 0.1 |
| GRAN (10<sup>3</sup>/μl) | 1.0 ± 0.4 | 1.0 ± 0.3 | 0.8 ± 0.4 | 1.2 ± 0.3 |
| LYM % | 85.3 ± 2.5 | 80.4 ± 3.6* | 86.7 ± 3.6 | 83.6 ± 1.8 |
| MONO % | 4.3 ± 0.2 | 5.5 ± 0.8** | 4.4 ± 1.2 | 5.1 ± 0.6* |
| GRAN % | 10.5 ± 2.4 | 13.6 ± 3.4 | 8.9 ± 2.6 | 11.3 ± 1.4 |
| HCT (%) | 44.4 ± 1.9 | 43.4 ± 2.2 | 45.5 ± 2.5 | 46.6 ± 2.1 |
| MCV (fl) | 44.1 ± 0.7 | 45.3 ± 1.3* | 45.4 ± 0.9** | 45.5 ± 0.5** |
| RDW<sub>a</sub> (fl) | 24.0 ± 0.6 | 25.6 ± 1.0** | 25.2 ± 0.6** | 26.3 ± 0.8*** |
| RDW% | 17.2 ± 0.5 | 18.0 ± 0.3** | 17.5 ± 0.6 | 18.9 ± 1.1** |
| HGB (g/dl) | 15.1 ± 0.6 | 14.7 ± 0.8 | 15.5 ± 0.7 | 15.9 ± 0.6* |
| MCHC (g/dl) | 34.0 ± 0.3 | 34.0 ± 0.4 | 34.1 ± 0.5 | 34.2 ± 0.6 |
| MCH (pg) | 15.0 ± 0.3 | 15.4 ± 0.4* | 15.5 ± 0.3** | 15.6 ± 0.2*** |
| RBC (10<sup>6</sup>/μl) | 10.1 ± 0.4 | 9.6 ± 0.7 | 10.0 ± 0.5 | 10.2 ± 0.4 |
| PLT (10<sup>3</sup>/μl) | 329.6 ± 85.8 | 320.5 ± 46.6 | 353.8 ± 80.4 | 314.2 ± 49.3 |
| MPV (fl) | 5.6 ± 0.4 | 5.5 ± 0.2 | 5.3 ± 0.2 | 5.7 ± 0.2 |

\*P ≤ 0.05; \*\*P ≤ 0.01; \*\*\*P ≤ 0.001; unpaired t-test relative to each cre control group. Numbers of mice are indicated in the table. WBC, white blood cells; LYM, lymphocytes; MONO, monocytes; GRAN, granulocytes; HCT, hematocrit; MCV, mean corpuscular volume; RDW, Red cell distribution width; HGB, hemoglobin; MCHC, mean corpuscular hemoglobin concentration; MCH, mean corpuscular hemoglobin; RBC, red blood cells.

**Table S5.**
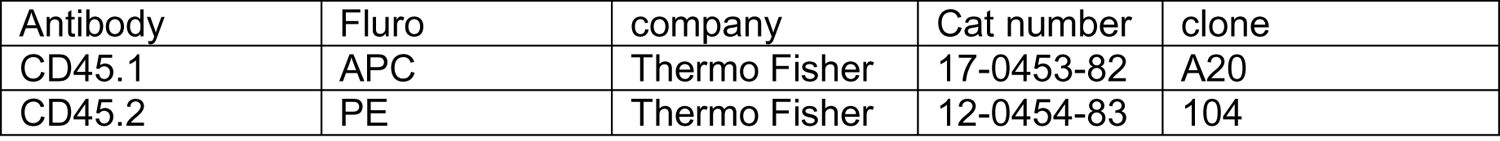
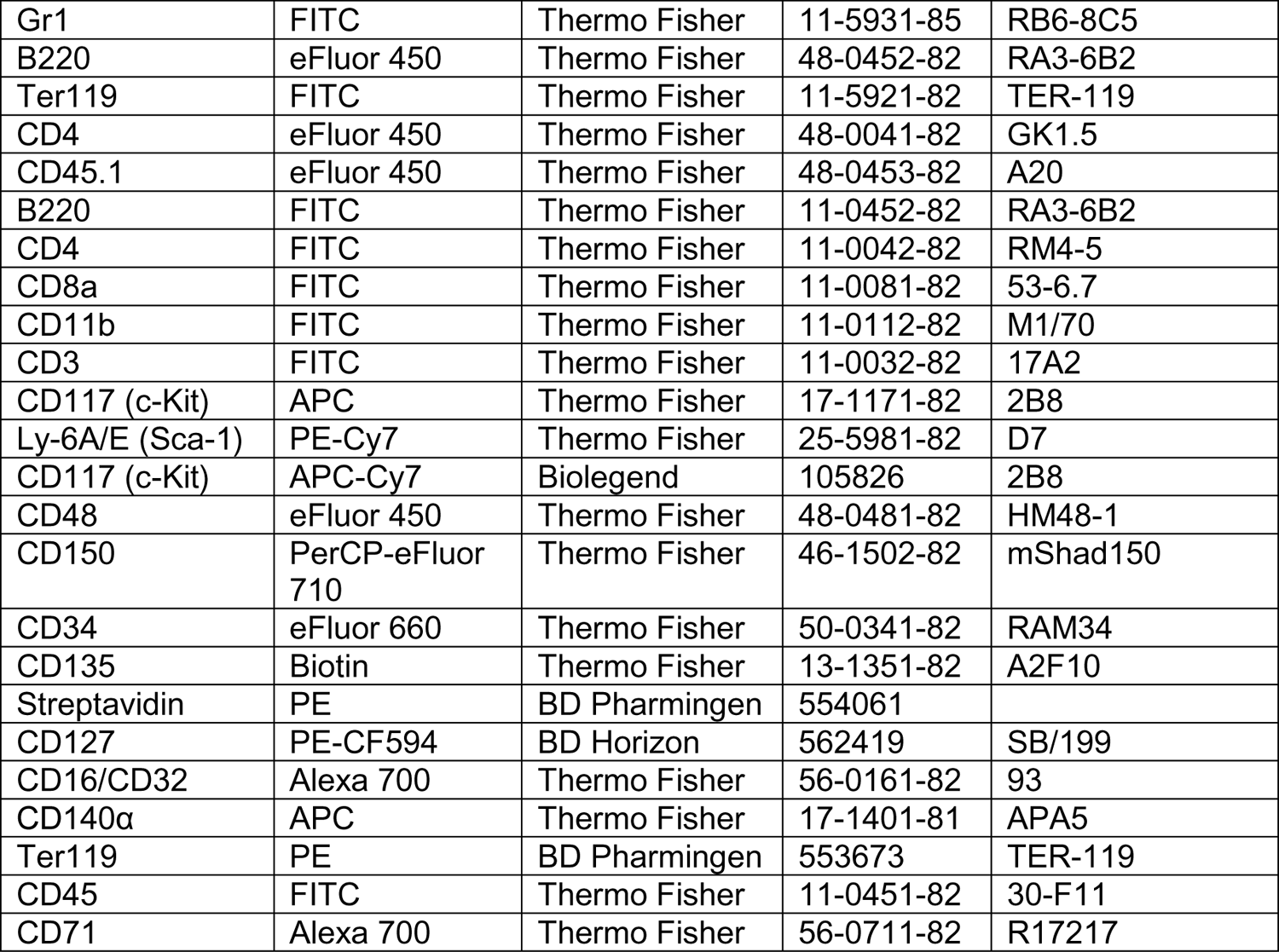
Antibodies used.

**Table S6.**
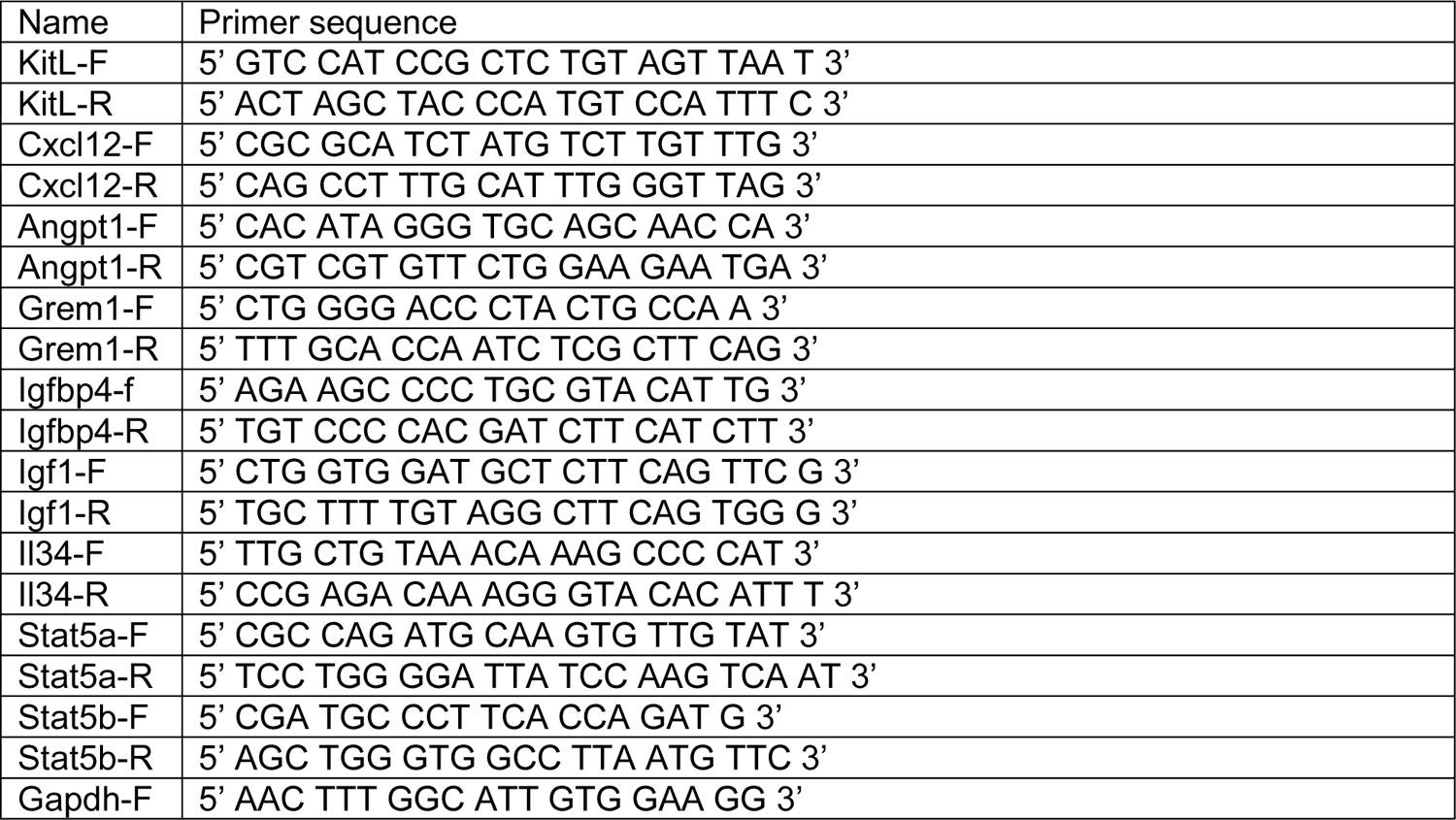
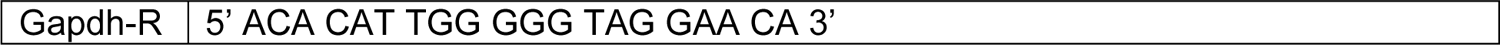
List of RT-PCR primers

## References

1. Yu VWC, Yusuf RZ, Oki T, Wu J, Saez B, Wang X, et al. Epigenetic Memory Underlies Cell-Autonomous Heterogeneous Behavior of Hematopoietic Stem Cells. Cell. 2017;168(5):944–5.

2. Grover A, Sanjuan-Pla A, Thongjuea S, Carrelha J, Giustacchini A, Gambardella A, et al. Single-cell RNA sequencing reveals molecular and functional platelet bias of aged haematopoietic stem cells. Nature communications. 2016;7:11075.

3. Sanjuan-Pla A, Macaulay IC, Jensen CT, Woll PS, Luis TC, Mead A, et al. Platelet-biased stem cells reside at the apex of the haematopoietic stem-cell hierarchy. Nature. 2013;502(7470):232 -6.

4. Pietras EM, Reynaud D, Kang YA, Carlin D, Calero-Nieto FJ, Leavitt AD, et al. Functionally Distinct Subsets of Lineage-Biased Multipotent Progenitors Control Blood Production in Normal and Regenerative Conditions. Cell stem cell. 2015;17(1):35–46.

5. Morita Y, Ema H, Nakauchi H. Heterogeneity and hierarchy within the most primitive hematopoietic stem cell compartment. JExpMed. 2010;207(6):1173-82.

6. Beerman I, Bhattacharya D, Zandi S, Sigvardsson M, Weissman IL, Bryder D, et al. Functionally distinct hematopoietic stem cells modulate hematopoietic lineage potential during aging by a mechanism of clonal expansion. ProcNatlAcadSciUSA. 2010;107(12):5465-70.

7. Zhou BO, Yu H, Yue R, Zhao Z, Rios JJ, Naveiras O, et al. Bone marrow adipocytes promote the regeneration of stem cells and haematopoiesis by secreting SCF. Nat Cell Biol. 2017;19(8):891–903.

8. Zhou BO, Yue R, Murphy MM, Peyer JG, Morrison SJ. Leptin-receptor-expressing mesenchymal stromal cells represent the main source of bone formed by adult bone marrow. Cell stem cell. 2014;15(2):154–68.

9. Asada N, Kunisaki Y, Pierce H, Wang Z, Fernandez NF, Birbrair A, et al. Differential cytokine contributions of perivascular haematopoietic stem cell niches. Nat Cell Biol. 2017;19(3):214–23.

10. Greenbaum A, Hsu YM, Day RB, Schuettpelz LG, Christopher MJ, Borgerding JN, et al. CXCL12 in early mesenchymal progenitors is required for haematopoietic stem-cell maintenance. Nature. 2013;495(7440):227-30.

11. Dahlin JS, Hamey FK, Pijuan-Sala B, Shepherd M, Lau WWY, Nestorowa S, et al. A single-cell hematopoietic landscape resolves 8 lineage trajectories and defects in Kit mutant mice. Blood. 2018;131(21):e1–e11.

12. Baryawno N, Przybylski D, Kowalczyk MS, Kfoury Y, Severe N, Gustafsson K, et al. A Cellular Taxonomy of the Bone Marrow Stroma in Homeostasis and Leukemia. Cell. 2019;177(7):1915–32.e16.

13. Tikhonova AN, Dolgalev I, Hu H, Sivaraj KK, Hoxha E, Cuesta-Dominguez A, et al. The bone marrow microenvironment at single-cell resolution. Nature. 2019;569(7755):222-8.

14. Schurch CM, Riether C, Ochsenbein AF. Cytotoxic CD8+ T cells stimulate hematopoietic progenitors by promoting cytokine release from bone marrow mesenchymal stromal cells. Cell stem cell. 2014;14(4):460–72.

15. Hirata Y, Furuhashi K, Ishii H, Li HW, Pinho S, Ding L, et al. CD150(high) Bone Marrow Tregs Maintain Hematopoietic Stem Cell Quiescence and Immune Privilege via Adenosine. Cell stem cell. 2018;22(3):445–53.e5.

16. Chow A, Lucas D, Hidalgo A, Mendez-Ferrer S, Hashimoto D, Scheiermann C, et al. Bone marrow CD169+ macrophages promote the retention of hematopoietic stem and progenitor cells in the mesenchymal stem cell niche. J Exp Med. 2011;208(2):261–71.

17. Pinho S, Marchand T, Yang E, Wei Q, Nerlov C, Frenette PS. Lineage-Biased Hematopoietic Stem Cells Are Regulated by Distinct Niches. Dev Cell. 2018;44(5):634–41.e4.

18. Cordeiro Gomes A, Hara T, Lim VY, Herndler-Brandstetter D, Nevius E, Sugiyama T, et al. Hematopoietic Stem Cell Niches Produce Lineage-Instructive Signals to Control Multipotent Progenitor Differentiation. Immunity. 2016;45(6):1219–31.

19. Yamashita M, Passegue E. TNF-alpha Coordinates Hematopoietic Stem Cell Survival and Myeloid Regeneration. Cell stem cell. 2019.

20. Golan K, Kumari A, Kollet O, Khatib-Massalha E, Subramaniam MD, Ferreira ZS, et al. Daily Onset of Light and Darkness Differentially Controls Hematopoietic Stem Cell Differentiation and Maintenance. Cell stem cell. 2018.

21. Wang Z, Li G, Tse W, Bunting KD. Conditional deletion of STAT5 in adult mouse hematopoietic stem cells causes loss of quiescence and permits efficient nonablative stem cell replacement. Blood. 2009;113(20):4856–65.

22. Lee J, Seong S, Kim JH, Kim K, Kim I, Jeong BC, et al. STAT5 is a key transcription factor for IL-3-mediated inhibition of RANKL-induced osteoclastogenesis. Scientific reports. 2016;6:30977.

23. Ogilvy S, Metcalf D, Gibson L, Bath ML, Harris AW, Adams JM. Promoter elements of vav drive transgene expression in vivo throughout the hematopoietic compartment. Blood. 1999;94(6):1855–63.

24. Stadtfeld M, Graf T. Assessing the role of hematopoietic plasticity for endothelial and hepatocyte development by non-invasive lineage tracing. Development. 2005;132(1):203–13.

25. Wang Z, Medrzycki M, Bunting ST, Bunting KD. Stat5-deficient hematopoiesis is permissive for Myc-induced B-cell leukemogenesis. Oncotarget. 2015;6(30):28961–72.

26. Qu P, Wang L, Min Y, McKennett L, Keller JR, Lin PC. Vav1 Regulates Mesenchymal Stem Cell Differentiation Decision Between Adipocyte and Chondrocyte via Sirt1. Stem Cells. 2016;34(7):1934–46.

27. Oguro H, Ding L, Morrison SJ. SLAM family markers resolve functionally distinct subpopulations of hematopoietic stem cells and multipotent progenitors. Cell stem cell. 2013;13(1):102–16.

28. Wilson A, Laurenti E, Oser G, van der Wath RC, Blanco-Bose W, Jaworski M, et al. Hematopoietic stem cells reversibly switch from dormancy to self-renewal during homeostasis and repair. Cell. 2008;135(6):1118–29.

29. Cabezas-Wallscheid N, Klimmeck D, Hansson J, Lipka DB, Reyes A, Wang Q, et al. Identification of regulatory networks in HSCs and their immediate progeny via integrated proteome, transcriptome, and DNA methylome analysis. Cell stem cell. 2014;15(4):507–22.

30. Miao R, Chun H, Feng X, Gomes AC, Choi J, Pereira JP. Competition between hematopoietic stem and progenitor cells controls hematopoietic stem cell compartment size. Nature communications. 2022;13(1):4611.

31. Boulais PE, Mizoguchi T, Zimmerman S, Nakahara F, Vivie J, Mar JC, et al. The Majority of CD45(-) Ter119(-) CD31(-) Bone Marrow Cell Fractio n Is of Hematopoietic Origin and Contains Erythroid and Lymphoid Progenitors. Immunity. 2018;49(4):627–39.e6.

32. Iacono G, Massoni-Badosa R, Heyn H. Single-cell transcriptomics unveils gene regulatory network plasticity. Genome biology. 2019;20(1):110.

33. van Dijk D, Sharma R, Nainys J, Yim K, Kathail P, Carr AJ, et al. Recovering Gene Interactions from Single-Cell Data Using Data Diffusion. Cell. 2018;174(3):716–29.e27.

34. Wang Z, Li G, Bunting KD. STAT5 N-domain deleted isoforms are naturally occurring hypomorphs partially rescued in hematopoiesis by transgenic Bcl-2 expression. Am J Blood Res. 2014;4(1):20–6.

35. Li G, Miskimen KL, Wang Z, Xie XY, Brenzovich J, Ryan JJ, et al. STAT5 requires the N-domain for suppression of miR15/16, induction of bcl-2, and survival signaling in myeloproliferative disease. Blood. 2010;115(7):1416–24.

36. Li G, Wang Z, Zhang Y, Kang Z, Haviernikova E, Cui Y, et al. STAT5 requires the N-domain to maintain hematopoietic stem cell repopulating function and appropriate lymphoid-myeloid lineage output. ExpHematol. 2007;35(11):1684–94.

37. Dai X, Chen Y, Di L, Podd A, Li G, Bunting KD, et al. Stat5 is essential for early B cell development but not for B cell maturation and function. JImmunol. 2007;179(2):1068–79.

38. Barnstein BO, Li G, Wang Z, Kennedy S, Chalfant C, Nakajima H, et al. Stat5 expression is required for IgE-mediated mast cell function. JImmunol. 2006;177(5):3421–6.

39. Couldrey C, Bradley HL, Bunting KD. A STAT5 modifier locus on murine chromosome 7 modulates engraftment of hematopoietic stem cells during steady-state hematopoiesis. Blood. 2005;105(4):1476–83.

40. Bradley HL, Couldrey C, Bunting KD. Hematopoietic-repopulating defects from STAT5-deficient bone marrow are not fully accounted for by loss of thrombopoietin responsiveness. Blood. 2004;103(8):2965–72.

41. Shelburne CP, McCoy ME, Piekorz R, Sexl V, Roh KH, Jacobs-Helber SM, et al. Stat5 expression is critical for mast cell development and survival. Blood. 2003;102(4):1290–7.

42. Bunting KD, Bradley HL, Hawley TS, Moriggl R, Sorrentino BP, Ihle JN. Reduced lymphomyeloid repopulating activity from adult bone marrow and fetal liver of mice lacking expression of STAT5. Blood. 2002;99(2):479–87.

43. Bradley HL, Hawley TS, Bunting KD. Cell intrinsic defects in cytokine responsiveness of STAT5-deficient hematopoietic stem cells. Blood. 2002;100(12):3983–9.

44. Snow JW, Abraham N, Ma MC, Goldsmith MA. Bone marrow transplant completely rescues hematolymphoid defects in STAT5A/5B-deficient mice. ExpHematol. 2003;31(12):1247–52.

45. Snow JW, Abraham N, Ma MC, Bronson SK, Goldsmith MA. Transgenic bcl-2 is not sufficient to rescue all hematolymphoid defects in STAT5A/5B-deficient mice. ExpHematol. 2003;31(12):1253-8.

46. Snow JW, Abraham N, Ma MC, Abbey NW, Herndier B, Goldsmith MA. STAT5 promotes multilineage hematolymphoid development in vivo through effects on early hematopoietic progenitor cells. Blood. 2002;99(1):95–101.

47. Yao Z, Cui Y, Watford WT, Bream JH, Yamaoka K, Hissong BD, et al. Stat5a/b are essential for normal lymphoid development and differentiation. ProcNatlAcadSciUSA. 2006;103(4):1000–5.

48. Berger A, Hoelbl-Kovacic A, Bourgeais J, Hoefling L, Warsch W, Grundschober E, et al. PAK-dependent STAT5 serine phosphorylation is required for BCR-ABL-induced leukemogenesis. Leukemia. 2014;28(3):629–41.

49. Hoelbl A, Schuster C, Kovacic B, Zhu B, Wickre M, Hoelzl MA, et al. Stat5 is indispensable for the maintenance of bcr/abl-positive leukaemia. EMBO MolMed. 2010;2(3):98–110.

50. Luc S, Anderson K, Kharazi S, Buza-Vidas N, Boiers C, Jensen CT, et al. Down-regulation of Mpl marks the transition to lymphoid-primed multipotent progenitors with gradual loss of granulocyte-monocyte potential. Blood. 2008;111(7):3424–34.

51. Drayer AL, Boer AK, Los EL, Esselink MT, Vellenga E. Stem cell factor synergistically enhances thrombopoietin-induced STAT5 signaling in megakaryocyte progenitors through JAK2 and Src kinase. Stem Cells. 2005;23(2):240–51.

52. Boer AK, Drayer AL, Rui H, Vellenga E. Prostaglandin-E2 enhances EPO-mediated STAT5 transcriptional activity by serine phosphorylation of CREB. Blood. 2002;100(2):467–73.

53. Kalaitzidis D, Neel BG. Flow-cytometric phosphoprotein analysis reveals agonist and temporal differences in responses of murine hematopoietic stem/progenitor cells. PLoSONE. 2008;3(11):e3776.

54. Ghanem S, Friedbichler K, Boudot C, Bourgeais J, Gouilleux-Gruart V, Regnier A, et al. STAT5A/5B-specific expansion and transformation of hematopoietic stem cells. Blood Cancer J. 2017;7(1):e514.

55. Kim Y, Lin Q, Glazer PM, Yun Z. Hypoxic tumor microenvironment and cancer cell differentiation. Curr Mol Med. 2009;9(4):425–34.

56. Rauch A, Haakonsson AK, Madsen JGS, Larsen M, Forss I, Madsen MR, et al. Osteogenesis depends on commissioning of a network of stem cell transcription factors that act as repressors of adipogenesis. Nature genetics. 2019;51(4):716–27.

57. Sehgal PB. Non-genomic STAT5-dependent effects at the endoplasmic reticulum and Golgi apparatus and STAT6-GFP in mitochondria. Jakstat. 2013;2(4):e24860.

58. Lee JE, Yang YM, Yuan H, Sehgal PB. Definitive evidence using enucleated cytoplasts for a nongenomic basis for the cystic change in endoplasmic reticulum structure caused by STAT5a/b siRNAs. Am J Physiol Cell Physiol. 2013;304(4):C312–23.

59. Lee JE, Yang YM, Liang FX, Gough DJ, Levy DE, Sehgal PB. Nongenomic STAT5-dependent effects on Golgi apparatus and endoplasmic reticulum structure and function. Am J Physiol Cell Physiol. 2012;302(5):C804–20.

60. Darvin P, Joung YH, Yang YM. JAK2-STAT5B pathway and osteoblast differentiation. Jakstat. 2013;2(4):e24931.

61. Mendez-Ferrer S, Lucas D, Battista M, Frenette PS. Haematopoietic stem cell release is regulated by circadian oscillations. Nature. 2008;452(7186):442-7.

62. Lucas D, Scheiermann C, Chow A, Kunisaki Y, Bruns I, Barrick C, et al. Chemotherapy-induced bone marrow nerve injury impairs hematopoietic regeneration. Nature medicine. 2013;19(6):695–703.

63. Hastings M, O’Neill JS, Maywood ES. Circadian clocks: regulators of endocrine and metabolic rhythms. J Endocrinol. 2007;195(2):187–98.

64. Marcheva B, Ramsey KM, Bass J. Circadian genes and insulin exocytosis. Cell Logist. 2011;1(1):32–6.

65. Nakao A. Temporal regulation of cytokines by the circadian clock. J Immunol Res. 2014;2014:614529.

66. Weger M, Diotel N, Dorsemans AC, Dickmeis T, Weger BD. Stem cells and the circadian clock. Dev Biol. 2017;431(2):111–23.

67. Chaudhari A, Gupta R, Patel S, Velingkaar N, Kondratov R. Cryptochromes regulate IGF-1 production and signaling through control of JAK2-dependent STAT5B phosphorylation. Molecular biology of the cell. 2017;28(6):834–42.

68. Grimley PM, Dong F, Rui H. Stat5a and Stat5b: fraternal twins of signal transduction and transcriptional activation. Cytokine Growth Factor Rev. 1999;10(2):131–57.

69. Gao P, Zhang Y, Liu Y, Chen J, Zong C, Yu C, et al. Signal transducer and activator of transcription 5B (STAT5B) modulates adipocyte differentiation via MOF. Cell Signal. 2015;27(12):2434–43.

70. Floyd ZE, Stephens JM. STAT5A promotes adipogenesis in nonprecursor cells and associates with the glucocorticoid receptor during adipocyte differentiation. Diabetes. 2003;52(2):308–14.

71. Yao Y, Bi Z, Wu R, Zhao Y, Liu Y, Liu Q, et al. METTL3 inhibits BMSC adipogenic differentiation by targeting the JAK1/STAT5/C/EBPβ pathway via an m(6)A-YTHDF2-dependent manner. FASEB journal: official publication of the Federation of American Societies for Experimental Biology. 2019;33(6):7529–44.

72. Hirose J, Masuda H, Tokuyama N, Omata Y, Matsumoto T, Yasui T, et al. Bone resorption is regulated by cell-autonomous negative feedback loop of Stat5-Dusp axis in the osteoclast. J Exp Med. 2014;211(1):153–63.

73. Seong S, Kim JH, Kim K, Kim I, Koh JT, Kim N. Alternative regulatory mechanism for the maintenance of bone homeostasis via STAT5-mediated regulation of the differentiation of BMSCs into adipocytes. Experimental & molecular medicine. 2021;53(5):848–63.

74. Grassinger J, Haylock DN, Williams B, Olsen GH, Nilsson SK. Phenotypically identical hemopoietic stem cells isolated from different regions of bone marrow have different biologic potential. Blood. 2010;116(17):3185–96.

75. Sacchetti B, Funari A, Michienzi S, Di CS, Piersanti S, Saggio I, et al. Self-renewing osteoprogenitors in bone marrow sinusoids can organize a hematopoietic microenvironment. Cell. 2007;131(2):324–36.

76. Morikawa S, Mabuchi Y, Kubota Y, Nagai Y, Niibe K, Hiratsu E, et al. Prospective identification, isolation, and systemic transplantation of multipotent mesenchymal stem cells in murine bone marrow. J Exp Med. 2009;206(11):2483–96.

77. Mendez-Ferrer S, Michurina TV, Ferraro F, Mazloom AR, Macarthur BD, Lira SA, et al. Mesenchymal and haematopoietic stem cells form a unique bone marrow niche. Nature. 2010;466(7308):829-34.

78. Soleimani M, Nadri S. A protocol for isolation and culture of mesenchymal stem cells from mouse bone marrow. Nature protocols. 2009;4(1):102–6.

79. Houlihan DD, Mabuchi Y, Morikawa S, Niibe K, Araki D, Suzuki S, et al. Isolation of mouse mesenchymal stem cells on the basis of expression of Sca-1 and PDGFR-alpha. Nature protocols. 2012;7(12):2103–11.

